# Powerful, efficient QTL mapping in *Drosophila melanogaster* using bulked phenotyping and pooled sequencing

**DOI:** 10.1101/2021.09.02.458801

**Authors:** Stuart J. Macdonald, Kristen M. Cloud-Richardson, Dylan J. Sims-West, Anthony D. Long

## Abstract

Despite the value of Recombinant Inbred Lines (RILs) for the dissection of complex traits, large panels can be difficult to maintain, distribute, and phenotype. An attractive alternative to RILs for many traits leverages selecting phenotypically-extreme individuals from a segregating population, and subjecting pools of selected and control individuals to sequencing. Under a bulked or extreme segregant analysis paradigm, genomic regions contributing to trait variation are revealed as frequency differences between pools. Here we describe such an extreme quantitative trait locus, or X-QTL mapping strategy that builds on an existing multiparental population, the DSPR (*Drosophila* Synthetic Population Resource), and involves phenotyping and genotyping a population derived by mixing hundreds of DSPR RILs. Simulations demonstrate that challenging, yet experimentally tractable X-QTL designs (>=4 replicates, >=5000 individuals/replicate, and a selection intensity of 5-10%) yield at least the same power as traditional RIL-based QTL mapping, and can localize variants with sub-centimorgan resolution. We empirically demonstrate the effectiveness of the approach using a 4-fold replicated X-QTL experiment that identifies 7 QTL for caffeine resistance. Two mapped X-QTL factors replicate loci previously identified in RILs, 6/7 are associated with excellent candidate genes, and RNAi knock-downs support the involvement of 4 genes in the genetic control of trait variation. For many traits of interest to drosophilists a bulked phenotyping/genotyping X-QTL design has considerable advantages.

## Introduction

There has been tremendous progress in understanding the genetic underpinnings of complex, polygenic trait variation in the last 20-30 years. This has been particularly true for complex human disease (Visscher *et al*. 2017), where the pioneering work of the Wellcome Trust Case-Control Consortium (Wellcome Trust Case Control Consortium 2007) led to an ever expanding catalog of thousands of replicable GWAS (genomewide association study) hits for a wide spectrum of human disorders and disease-relevant traits (Buniello *et al*. 2019). Equally, the efficiency and power of studies dissecting complex trait variation in animal and plant systems has been enhanced by inexpensive, sequencing-based genotyping solutions (e.g., (Davies *et al*. 2016)), and an array of readily available genetic mapping populations (e.g., (Kover *et al*. 2009; Aylor *et al*. 2011; Svenson *et al*. 2012; King *et al*. 2012b; Rat Genome Sequencing and Mapping Consortium *et al*. 2013; Huang *et al*. 2014; Noble *et al*. 2021)).

Whereas GWAS is the dominant mapping paradigm in humans, the experimental flexibility of animal and plant systems - particularly the traditional “model systems” (e.g., *Drosophila melanogaster*, *Caenorhabditis elegans*, mice) - has facilitated the development of an array of mapping approaches with varied strengths and weaknesses. GWAS-based methods, such as the DGRP (*Drosophila* Genetic Reference Panel; (Huang *et al*. 2014)), a set of 200 inbred lines derived from wild-caught flies, can have high power to identify intermediate-frequency loci of large effect (e.g. (Magwire *et al*. 2012)), and exhibit fine mapping resolution given that they leverage the extensive ancestral recombination experienced by the sampled population. But as with all GWAS approaches such methods will struggle when functional alleles are rare (Pritchard 2001; Spencer *et al*. 2009; Thornton *et al*. 2013), or variant effects are low (Long *et al*. 2014; Mackay and Huang 2018). Other strategies in experimental plants and animals use multiparental populations (MPPs), collections of recombinant individuals or inbred lines derived from intercrossing a modest number of founding genotypes for a number of generations. One example of a *D. melanogaster* MPP is the DSPR (*Drosophila* Synthetic Population Resource; (King *et al*. 2012a; b)), which consists of two sets of RILs, each derived from an independent set of 8 founding inbred strains. MPP-based methods sample less population diversity than GWAS designs, yet can have high QTL (Quantitative Trait Loci) mapping power (King *et al*. 2012a; Gatti *et al*. 2014; Keele *et al*. 2019). This is true even when causative variants are rare in the population, since those captured in the founders are present at higher frequency in the mapping panel. Additionally, the design and analysis of MPPs explicitly enables the detection of loci segregating for multiple functional alleles (Long *et al*. 2014); When directly assessed, such allelic heterogeneity appears to be common for complex traits (King *et al*. 2014; Hormozdiari *et al*. 2017).

Despite the range of tools available, a complete empirical picture of the genetic basis of complex traits is lacking. One practical barrier to the use of available mapping panels in the *D. melanogaster* system is that power is a strong function of sample size, so hundreds of strains must be maintained, and multiple individuals must be phenotyped from each strain. Methods that mitigate some of the burden of phenotyping can be enabling.

One attractive alternative to depending on inbred lines is to employ a mapping approach that leverages a bulked phenotyping strategy (Figure 1), akin to the X-QTL approach used by yeast geneticists (Ehrenreich *et al*. 2012). An X-QTL strategy entails creating a large “base” population of recombinant individuals, and subjecting a sample to a bulked phenotyping regime that selects the top (say) 5-10% of individuals. This selection step enriches for haplotypes that harbor phenotype-increasing alleles closely-linked to a causative gene. Haplotype frequencies are then estimated in pooled DNA samples obtained from the base and selected populations, and at sliding windows throughout the genome a statistical test identifies regions showing significant divergence between the populations. In contrast to mapping using an individual- or RIL-based strategy, where single individuals or sets of genetically-identical individuals are phenotyped, an X-QTL approach is attractive: First, an investigator need only handle/maintain/track a single population as opposed to several hundred separate RILs. Second, deriving a selected population has the potential to be straightforward, especially for traits that “phenotype themselves” - e.g., survival or preference traits. Third, phenotyped individuals are outbred diploids raised in a common garden environment, so any effects of inbreeding depression are alleviated, and batch/vial effects are mitigated. Fourth, the sequencing effort required to accurately estimate haplotype frequencies from pooled samples initiated via a small number of founders (e.g., 8) is - perhaps counter-intuitively - modest (explored below).

**Figure 1:**
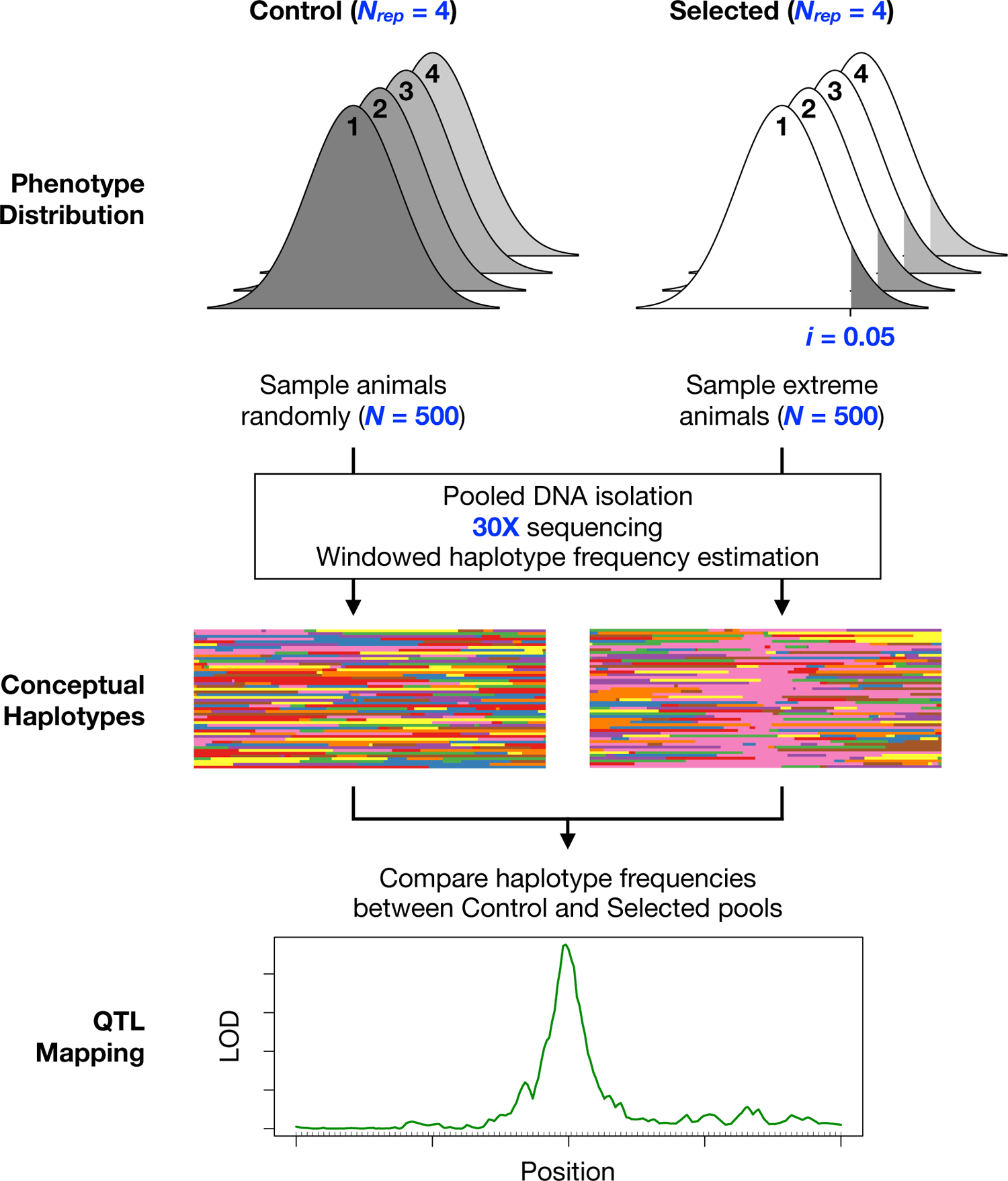
Overview of the DSPR X-QTL approach. A segregating base population is created by mixing a series of DSPR RILs. For each of several replicates (*N_rep_*), a set of *N_DNA_* individuals are randomly sampled from the base population (control pool), along with a set of *N_DNA_* phenotypically-extreme individuals (selected pool) in - for instance - the top 5% of the phenotype distribution (*i*). Following DNA extraction, each pool is sequenced to ∼30X coverage, and haplotypes are estimated in windows through the genome. Haplotype frequencies are then compared between paired control and selected pools, and QTL are evident as significant differences in frequency.

Here, we consider an X-QTL experiment utilizing the set of DSPR RILs (King *et al*. 2012b). Our design consists of intercrossing a set of RILs (*N_RIL_*) to create a base population, and expanding this population for several generations to both grow the total number of flies available for testing, and further recombine the RILs utilized (Figure 1). We then impose a selective regime on the base population, select for the *i*% most extreme individuals, obtain pooled DNA preps (of size *N_DNA_*) from both the base and selected samples, sequence, and repeat the entire experiment *N_rep_* times. Based on the observed frequencies of SNPs throughout the genome, and the known sequence of the 8 highly-characterized inbred DSPR founders (King *et al*. 2012b; Chakraborty *et al*. 2019), a vector of 8 haplotype frequencies can be estimated at locations throughout the genome for each pooled sample. Finally, at those same locations we carry out a statistical test for differentiation between the selected and control populations; loci associated with large values of the test statistic indicate the presence of a QTL. The X-QTL strategy we propose here is analogous to earlier work (Ehrenreich *et al*. 2012; Burga *et al*. 2019), but differs in having multiple founder alleles segregating in the base population, and in establishing the base from RILs derived from an advanced generation intercross in order to obtain higher mapping resolution. Below we explore - in theory and in practice - the utility of this approach relative to typical RIL-based mapping as a function of achievable experimental designs.

## Materials & Methods

### The *Drosophila* Synthetic Population Resource

The DSPR consists of a series of RILs derived from a pair of advanced generation synthetic, recombinant populations (populations A and B, or pA and pB), each of which was founded by a set of 8 highly-inbred founder strains (King *et al*. 2012b; Chakraborty *et al*. 2019). The synthetic populations were each maintained as duplicate subpopulations (pA1, pA2, pB1, pB2) at large census size for 50 generations of free recombination, after which RILs were derived with 25 generations of strict brother-sister mating. The resulting RILs are highly inbred (King *et al*. 2012b).

### Power simulations

To assess the power of the X-QTL approach in comparison with RIL-based mapping using the DSPR, we simulated the distribution of LOD scores for a complex trait with 50% heritability, 2% heritability due to a focal gene, and variation at this gene due to 8 equally-frequent founder haplotypes with additive Gaussian genotypic effects. These assumptions are reasonable since QTL exhibiting >2% heritability are commonly observed in RIL-based QTL mapping studies employing the DSPR (see for instance (Najarro *et al*. 2015)and(Everman *et al*. 2019)), Beavis effects are subtle for large experiments (Beavis *et al*. 1991; Beavis 1994; King and Long 2017), and QTL mapped in the DSPR often appear to exhibit an allelic series (King *et al*. 2014). Changing the trait heritability while holding the heritability of the focal gene constant should not impact the power of the X-QTL approach. But for RIL-based mapping, replicate phenotypic measures for each strain will increase power, since the ratio of environmental variation to non-focal locus genetic variation is increased. To estimate power at a marker located in a single focal QTL we need only simulate the QTL itself and can ignore any flanking information.

For each of 1,000 replicates we simulate a RIL-based mapping experiment using 500 randomly-chosen RILs from a single population, 10 phenotype measures per RIL, and derive genomewide *P*-values via one-way ANOVAs with founder haplotype as a factor. To simulate the X-QTL experiment we create 4, 8, or 12 large replicate (*N_rep_*) diploid base populations by mixing the above 500 RILs, instantaneously expand the population, then choose the 5% or 10% most phenotypically-extreme individuals from each replicate population to create selected pools. For each replicate we randomly choose 150, 300, or 600 diploid individuals from the selected pool, and an equivalent number from the expanded base population to create a control pool (treatment = *trt*). The 8 *known* founder haplotype (*h*) arcsine square-root transformed frequencies (*asf*) at each position are tested for differentiation between control and selected pools using an ANOVA (i.e., *asf*∼*h*trt*, with *h:trt* tested using *replicate:h:trt*). Minus log base 10 *P*-values, −log_10_(*P*), associated with the X-QTL and RIL-based statistical tests are compared.

### False positive rate and significance thresholds

We sampled (*N_RIL_*) 200, 400, 600, or 800 DSPR pA RILs, extracting haplotype probabilities every 200-kb along chromosome 3L. At each location haplotype calls are represented by 8 additive dose probabilities (i.e., the vector sums to 1), but here we more simply define the founder haplotype at each position in each RIL as the largest of the eight dosage calls. Since doses are typically near 0 or 1 (King *et al*. 2012b) switching from soft to hard calls does not discard a great deal of information. Sampling only every 200-kb (as opposed to the native 10-kb marker spacing in the DSPR) makes the simulation machinery efficient, and since adjacent test statistics are highly correlated at sub-cM sampling densities there is little need to sample at a higher marker density (see Figure S2A). We simulated a Gaussian QTL in the middle of the chromosome contributing 2% to heritable variation. Additionally, we placed additive Gaussian polygenic “background” loci of equal variance at each of the 200 marker locations such that the total background variance was 8%, and environmental variation was set equal to 90% (*V_e_* in this manner includes genetic variation due to the remainder of the genome as well as variance due purely to the environment). Unlike for power calculations (above), after expansion of the base population to 20 thousand individuals we allow 4, 8, or 16 generations of random mating, with exactly one recombination event per female chromosome per generation.

For each starting number of RILs (*N_RIL_*), we simulated selection intensities of 5, 10, and 20% (*i*) and randomly sampled two sets of 150, 300, or 600 individuals (*N_DNA_*) from a set of 20,000 simulated individuals (i.e., mock treatment vs. control) to make DNA. Errors in the haplotype estimator and/or differences in the number of DNA molecules per individual that make their way into the sequencing library, are likely to reduce the number of individuals being tested. Thus, we consider these values of *N_DNA_* to reflect the idea of *an effective sample size* that is lower than the actual number of individuals that would be used in an experiment (i.e., *N_DNA_* values of 150/300/600 might represent census sizes of perhaps 250/500/1000). We repeated the simulation 4, 8, or 12 times for each parameter combination mimicking an experiment replicated to different levels, with each experiment producing the same minus log 10 *P*-value as the power calculations described above. For each parameter set we calculate the per chromosome false positive rate (FPR) as the proportion of scans out of 250 pure replicates in which a single marker has a LOD score greater than 4. Similarly, we calculate the genomewide false positive rate as gFPR = 1−(1−FPR)^5^, since our test chromosome 3L is roughly 1/5^th^ of the *D. melanogaster* genome. At a LOD threshold of 4 the gFPR is 7.1% (i.e., 7.1% of all genome scans will result in a single interval with a LOD > 4). But the false positive rate is a subtle function of the manipulated parameters, and the number of flies contributing to DNA pools has a measurable impact on the gFPR. Conditioning on effective pools of 300 or 600 flies, mimicking census sizes of perhaps 500 or 1000, results in an gFPR of 3.9%. Thus a LOD score of 4 seems to be an appropriate threshold over a large range of parameters, but smaller pools may result in a slightly liberal false positive rate.

### X-QTL localization and false positives due to polygenic background

In the simulation described above we estimated localization ability in kilobases as the number of markers within a 2-LOD drop of the most-significant marker (MSM) in a scan, conditional on the MSM being greater than 4, multiplied by 200 (since markers are spaced every 200-kb). We further examined the distribution of LOD scores at markers greater than 7-Mb from the causative site; Elevated LOD scores at such loosely-linked sites likely reflect signal due to linkage disequilibrium between founder haplotypes at the locus and chromosome-wide polygenic scores. For some of the above parameter combinations the localization ability was approaching the marker density of one every 200-kb. As a result we repeated the localization experiment for a subset of the parameter combinations, but changed the marker density to one every 10-kb, and only examined the middle third of chromosome 3L. We further only considered four generations of random mating following derivation of the base population. These two changes held the considerable computation effort roughly constant, and allowed us to more precisely examine localization ability. We then estimated the localization ability in kilobases as ten times the number of markers (since markers are spaced every 10-kb) within a two LOD drop of the MSM in a scan, conditional on the MSM being greater than 4. We carried out ANOVAs on the average interval size as a function of the parameters we varied (*N_RIL_*, *i*, *N_DNA_*, *N_rep_*) and included the average LOD score per parameter combination as a covariate.

### Development of a mixed DSPR population

To create an experimental mixed population of RILs we manually collected 10 eggs from each of 663 pA RILs (365 from subpopulation pA1, 298 from subpopulation pA2), placing ∼400 eggs into each of a series of *Drosophila* stock bottles (6 oz; ThermoFisher, AS355) containing ∼50ml of cornmeal-yeast-molasses medium. All bottles were placed inside a 1 cubic foot (12-inch × 12-inch × 12-inch) population cage, and approximately every 2 weeks old bottles were removed from the cage and replaced with 9-12 fresh bottles. The number of flies in the cage remained very high throughout the execution of the experiments described below.

### Rearing and collecting experimental animals

Four replicates of the experiment were carried out, following 1 (replicate A), 3 (rep. B), 4 (rep. C) and 5 (rep. D) generations of population cage maintenance. In each case we followed the same basic procedure outlined below (also see Table S1).

**Table 1:** Properties of mapped QTL

| QTL | Chr | Pos <sup>a</sup> | LOD | <u>2 LOD support</u> |  | Size (kb) | Candidate gene(s) | # Protein coding genes |
| --- | --- | --- | --- | --- | --- | --- | --- | --- |
|  |  |  |  | left <sup>a</sup> | right <sup>a</sup> |  |  |  |
| A | X | 3716075 | 5.36 | 3216075 | 4096075 | 880 | *Tlk* | 51 |
| B | 2L | 3370610 | 4.72 | 3110610 | 3890610 | 780 | *E23, Ugt36A1* | 105 |
| C | 2L | 11730610 | 4.59 | 11130610 | 12050610 | 920 | *Vha100-5, Crys* | 95 |
| D | 2R | 10848099 | 13.1 | 10688099 | 11128099 | 440 | *Cyp12d1* | 41 |
| E | 3L | 13108478 | 4.99 | 12508478 | 15708478 | 3200 | - | 315 |
| F | 3R | 14067353 | 8.81 | 13627353 | 14247353 | 620 | *Cyp6d5* | 79 |
| G | 3R | 26287353 | 6.77 | 25787353 | 26707353 | 920 | *osy* | 99 |

<sup>a</sup> Physical positions given with respect to Release 6 of the *D. melanogaster* reference genome.

We placed 5-7 apple juice agar dishes (Text S1) supplemented with a small quantity of live yeast paste (Text S1) into the population cage for several hours. Afterwards, any yeast paste was removed from the surface of the agar, the dishes were filled with 1X PBS (phosphate-buffer saline), and a paintbrush was used to dislodge the eggs from the surface. Subsequently the egg suspension from all plates was poured into a 50-ml tube and the eggs allowed to fall to the bottom. The majority of the PBS was poured off and replaced 2-3 times until the PBS was relatively clear and free of debris. Subsequently a standard 100-μl pipettor was set to 12-μl, and a wide-bore tip used to move aliquots of the egg suspension to rearing vials (narrow, polystyrene; ThermoFisher, AS515) containing ∼10ml of cornmeal-yeast-molasses media (Text S1). (Pipetting very aggressively ensures eggs are moved along with PBS). For each replicate we collected eggs over 2 consecutive days to generate a sufficient number of vials/experimental females. While we do not know the exact number of eggs per vial, the level of larval activity across vials was visually consistent, and in a sample of 10 rearing vials for which all emerging females were counted, the number of adult females averaged ∼70 per vial.

Following development, and 2 days after the first evidence of adults, all flies from each rearing vial were tipped to a mixed-sex housing vial. Given the density of flies in these vials they were placed on their side. Two days later, experimental female flies were manually collected over CO_2_ anesthesia and sorted into single-sex housing vials in groups of 19-30.

### Caffeine resistance assay

The assay used largely replicated that employed previously to identify QTL in the set of inbred DSPR RILs (Najarro *et al*. 2015). For each of the four replicates (A, B, C, and D) we carried out the following procedure.

The day prior to the initiation of the experiment we generated assay media (Text S1), cooled 1-liter to ∼55°C, and thoroughly mixed in 10-g of caffeine (SigmaAldrich, C0750) achieving a final caffeine concentration of 1%. Media was then poured into a series of 100-mm petri dishes and allowed to cool. Subsequently, bundles of polycarbonate *Drosophila* activity monitor tubes (5-mm diameter × 65-mm length; TriKinetics, PPT5×65) were filled to ∼10-mm height with solidified media by pressing them into the media surface, wiped clean, then sealed by plunging the tubes individually into molten paraffin wax.

On the day of the experiment, 3-5 day old females (*N* = 2,337-2,572 per replicate) were moved from single-sex housing vials and loaded into monitor tubes, and tubes were capped with small foam plugs cut from Droso-Plugs (Genesee Scientific, 59-200). Rather the placing the monitor tubes into an automated monitoring system (Najarro *et al*. 2015), tubes were placed on their side along the base of a series of cardboard/plastic narrow fly vial trays (Genesee Scientific, 32-124 or 59-163B), with ∼160 tubes per tray. Starting the day following setup, the tubes were surveyed for dead animals twice each day, and dead animals were counted and removed.

For the first replicate (replicate A), 43% of the animals were loaded into monitor tubes via manual, oral aspirators, while 57% were loaded via an automated instrument (FlySorter). By segregating tubes into different trays based on loading method, we noticed that automatically-loaded animals had lower lifespan on average (Figure S6). Since we selected the most resistant animals on a tray-by-tray basis (see below), the impact of the loading method may be limited (indeed, dropping replicate A from the final X-QTL analysis has little effect over dropping replicate B - Figure S4). However, for all three later replicates (replicates B, C, D) we solely loaded flies by manual aspiration.

### Collection of pools of control and caffeine-resistant animals

For each replicate we collected 250 “control” females from the set of single-sex housing vials by randomly selecting 25 vials, aspirating 10 females per vial into a 15-ml tube on ice, then freezing the tube at −20°C to await DNA isolation.

Ideally the pool of “selected” females for each replicate would be the 250 females surviving the longest under caffeine conditions. This was not possible for two reasons. First, we intermittently monitored the number of dead animals, and had to judge the appropriate time to collect the final set of females to avoid collecting many more, or many fewer, than the 250 desired. Second, the exposure cannot start for all experimental females at the same time; with 2-3 investigators it took 6-9 hours to load all females into tubes. Since our target was the 250 most resistant animals (i.e., 10% of the starting sample), and since tubes were sequentially arrayed out into trays at ∼160/tray, we targeted the collection of ∼16 females per tray. The mortality trajectory of each tray was fairly consistent (Figure S6), aside from trays of flies that were automatically loaded (see above). When we reached the target number of live flies per tray (subject to the intermittent monitoring constraint noted), those monitor tubes were frozen at −20°C, and subsequently all selected flies were pooled into a single 15-ml tube, and re-frozen at −20°C. We collected 228-254 females per selected pool (9.0-10.3% of the starting set of flies.).

### DNA isolation

DNA was isolated from each of the 8 samples of flies using an adaptation of the Gentra Puregene Cell Kit protocol (Qiagen, 158767). Briefly, each pool of flies was homogenized via glass beads in 1X PBS, and subjected to several strokes of a dounce homogenizer. Then a small amount of the homogenate was taken forward through cell lysis, protein precipitation, an RNase step, DNA precipitation and resuspension (see Text S2 for more details). DNA integrity was confirmed by running a small amount of each sample on a 1.5% agarose gel, and DNA was quantified using a fluorometer (Qubit dsDNA BR Assay Kit, ThermoFisher, Q32853).

### Library preparation and sequencing

Each DNA sample was diluted, and a 50-μl aliquot containing between 832-841 ng was provide to the KU Genome Sequencing Core for library construction using the NEBNext Ultra II DNA Library Prep Kit (NEB, E7645L), incorporating unique dual-indexing (NEB, E6440S). Final library sizes ranged from 303-bp to 327-bp (Agilent TapeStation 2200), and insert sizes are expected to be around 200-bp. Libraries were pooled at equal concentrations, and sequenced with PE150 reads on a single S4 lane of an Illumina NovaSeq 6000 (UC Irvine Genomics High-Throughput Facility).

### Haplotype estimation from experimental sequencing data

Haplotypes are estimated using slight modifications to published software (Linder *et al*. 2020). In short, we use `bwa-mem` (Li 2013) to generate a BAM file for each sequenced pooled sample, and for each of the highly inbred founder lines from which the pA DSPR population was derived (King *et al*. 2012b). We then use `bcftools mpileup` and `bcftools call` ((Li 2011); bcftools version 1.9) to generate a file that reports REF and ALT counts at all SNPs. Finally, we employ `bcftools query`, along with a custom perl script, to output the frequency of the REF allele in each sample for the set of SNPs that are not polymorphic within any given founder strain.

For each pooled sample, for a window size of 200kb and a step size of 10kb, we use the `limSolve` R library (Soetaert *et al*. 2009) to estimate the optimal vector of 8 founder haplotype frequencies (***f***) that minimizes the sum of the squares between observed allele frequencies and predicted allele frequencies; In other words, ***Y*** = ***Xf***, where ***Y*** is a vector of allele frequencies in the pool, and ***X*** is an *F* column matrix of (largely 0,1) genotypes associated with the *F* founders. The `limSolve` library allows us to constrain solutions to those whose frequencies sum to 1, and where each frequency is >0.03% (this small mass for every founder avoids convergence issues at 0 and 1). For any given window the number of SNP positions is large, and dropping subsets of SNPs does not impact the estimator very strongly. The script to do this is described more fully in prior work (Linder *et al*. 2020) and on GitHub.

### Establishing the level of error in haplotype estimation

We created a pseudo-pool with an average sequencing coverage of 240X by combining an equal number of reads from the replicate A and B caffeine experiment control pools (*N_DNA_* = 250 in each case). Using this high coverage data we estimated allele frequencies at a genomewide set of SNPs each private to a single founder. We then down-sampled the number of reads in the pool to more modest coverage (70X, 35X, 17X) and ran our haplotype frequency estimator (above). The frequency of a private SNP at 240X coverage, and the frequency of the founder harboring that private SNP at the window nearest the SNP location, are both estimators of the frequency of that founder in the sample. We calculated the mean square difference between the two estimators as a function of coverage as: 0.000708 (full 240X coverage), 0.000711 (70X), 0.000717 (35X), and 0.000725 (17X). The mean squared difference between frequency estimates (Var_diff_) should equal the variance in SNP estimating frequency (Var_SNP_) plus the variance contributed by the haplotype estimator (Var_HAP_). Since Var_diff_ varies little as a function of coverage, this suggests error in the haplotype frequency estimator is quite small.

We estimate the expectation of Var_SNP_ as *<u>pq</u>/<u>N</u>* over all SNPs at 240X to be 0.000322. But if we alternatively obtain the expectation of Var_SNP_ as the variance among SNP frequencies within the same founder allele and within 10-kb of one another (such SNPs should have almost identical frequency), we obtain 0.000611, a variance roughly twice as large. We conclude that the error in SNP frequency estimates obtained from a pooled DNA sample has an over-dispersed variance relative to binomial expectations, as is commonly claimed in the literature (King *et al*. 2012b; Wei *et al*. 2017; Zhang and Emerson 2019). If we assume 0.000611 more accurately estimates Var_SNP_, this implies the variance in our haplotype estimator is quite small, with an average absolute error of ∼0.01 (i.e., sqrt(Var_diff_-Var_SNP_)) irrespective of coverage.

### Functional testing of potential candidates

We used the Gal4-UAS-RNAi system to functionally test a series of genes implicated by mapped X-QTL. We used the ubiquitous, *Actin 5C* (*Act5C*) promoter-driven Gal4 strain (Bloomington Drosophila Stock Center number 25374), and a strain expressing Gal4 in the adult anterior midgut (1099 from Nicholas Buchon, flygut.epfl.ch; (Buchon *et al*. 2013)). All UAS-RNAi strains were from the VDRC (Vienna Drosophila Resource Center) “GD” collection, which each harbor a *P*-element-derived UAS transgene (Dietzl *et al*. 2007). Transgenes targeted genes *Crys* (stock ID 37736), *Cyp12d1* (50507), *Cyp6d5* (12138), *Ugt36A1* (9489), *E23* (2620), *osy* (38661), *Tlk* (46424), and *Vha100-5* (6121). We compared each to a background, transgene-free control strain (stock ID 60000).

We tested the same set of experimental Gal4-UAS genotypes across two batches. In the first batch we crossed ∼7 male Gal4 animals to 10 female UAS (or control) animals, setting up 2 replicate vials per Gal4/UAS combination (18 genotypes total). Two days following the first emergence of adults, all flies from each vial were tipped to a mixed-sex housing vial. After a further 2 days, experimental females were sorted over CO_2_ into single-sex vials housing 20 experimental females. The following day 32 experimental, 3-5 day old females (16 per replicate cross vial) were loaded without anesthesia into 1% caffeine monitor tubes (see above), and we used the *Drosophila* activity monitoring system to track movement of each individual, yielding an accurate lifespan for each (see (Najarro *et al*. 2015)).

In the second batch we crossed 10 female Gal4 animals to ∼7 male UAS (or control) animals - i.e., the reciprocal cross direction from the first batch. We set up 3 replicate vials per Gal4/UAS combination (16 genotypes total; 2 crosses involving the *Act5C*-Gal4 strain were not re-attempted since in the first batch no experimental Gal4-UAS-RNAi animals were obtained, Table S2). As described for the first batch we collected 3-5 day old experimental females, and loaded 48 animals per genotype (16 per cross vial) into monitors. Caffeine resistance phenotypes for each Gal4-UAS-RNAi genotype and each batch were compared to the relevant control genotype via Dunnett’s tests.

### Environmental conditions used to rear and assay X-QTL and RNAi flies

The source X-QTL population cage, the vials used to rear and house experimental X-QTL and RNAi animals, and the monitor tubes used to assay caffeine resistance were all resident in the same laboratory incubator, and maintained at 25°C, 50% relative humidity, on a 12 hour light: 12 hour dark cycle. Media used in population cage bottles, and in both rearing and housing vials was a cornmeal-yeast-molasses mix (Text S1), while media used for the resistance assay was a cornmeal-yeast-dextose mix supplemented with caffeine (Text S1).

### Data and analytical code availability

Simulation, analysis code, and code to reproduce many of the figures is available from GitHub (https://github.com/tdlong/fly_XQTL). Short-read sequencing data for all 8 control and caffeine-selected pools can be obtained from the NCBI SRA under bioproject accession PRJNA714149. RNAi data and R analysis scripts (rnai_supp_data_analysis.zip) and the haplotype calls for each population (allhaps.200kb.txt.gz) are available at FigShare (data uploaded to Figshare). The haplotype calls allow the mapping results to be reproduced without the computation effort associated with aligning raw reads to the genome, calling SNPs, and calling haplotypes.

## RESULTS

### Powerful DSPR-based X-QTL mapping in *Drosophila*

We simulated QTL mapping power with a standard RIL-based QTL mapping experiment, as well several different X-QTL designs. Figure 2 depicts the distribution of LOD scores at a QTL contributing 2% to trait variation. The distribution of the X-QTL LOD scores is presented as a function of the number of individuals sampled to create the selected and control DNA pools (150, 300, 600), the selection intensity applied during phenotyping (10% and 5%), and the number of replicate base populations (4, 8, 12). We see the expected null distribution of X-QTL test statistics when comparing two equally-sized draws from the control/base population (Figure S1), and provided DNA is prepared from a sufficiently large pool of animals, a LOD threshold of 4 holds the genomewide false positive rate at <=5%. Thus in Figure 2 power is the proportion of LOD scores greater than 4. Power with the X-QTL framework is routinely higher than using RILs directly. And it is clear that stronger selection, more replicates, and a greater number of individuals contributing to the DNA pools strongly impact X-QTL power (Figure 2), and should be maximized in any experiment.

**Figure 2:**
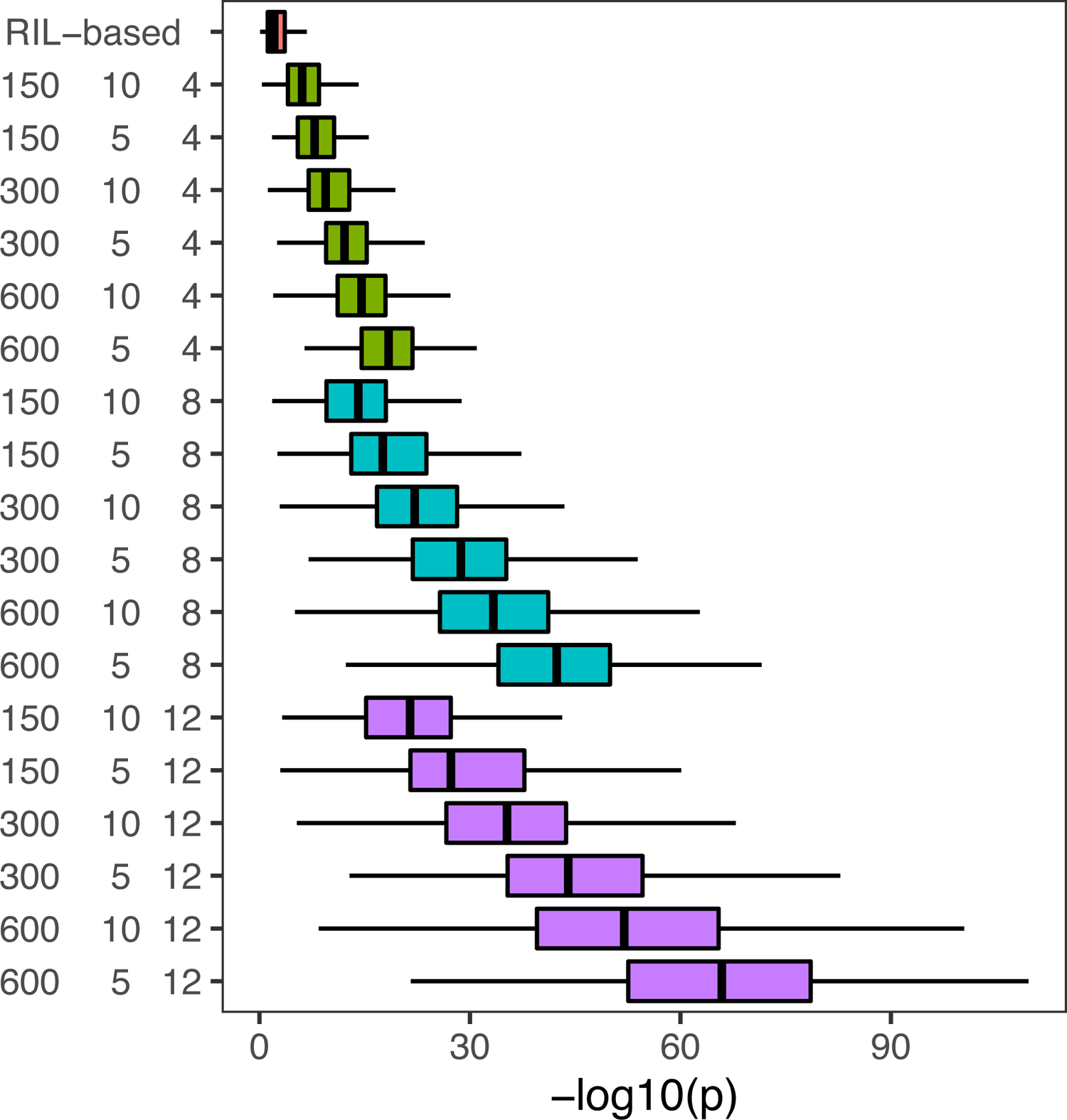
Distribution of −log_10_(*P*) test statistics for RIL-based mapping compared to X-QTL. Analysis assumes a QTL contributing 2% to heritability. X-QTL −log10(*p*-value) distributions are shown as a function of the effective number of individuals contributing to the DNA pools (150, 300, 600), selection intensity (10%, 5%), and the number of experimental replicates (4, 8, 12). Power is routinely higher via X-QTL mapping compared to mapping directly with RILs, and larger pools, more extreme selection, and a greater number of replicates, further increases X-QTL power.

The simulated pool sizes of 150, 300, or 600 individuals sampled to make genomic DNA should be thought of as the *effective number of individuals* in the pool; Factors such as uneven representation of individuals in the pool (e.g., variation in body size, differential lysis or degradation during the bulk DNA prep), and errors in haplotype frequency estimation will result in the effective number of individuals in the pool being lower than the census number. This being said, by only choosing flies of a single sex (to somewhat control for body size variation), using fresh tissue from living animals, and generating 20-40X sequencing coverage per pool (see experimental results below), we believe the difference between the effective and census population sizes can be minimized. We suggest that effective sizes of 150, 300, and 600 may approximate census sizes of 250, 500, and 1000 individuals, respectively. Overall, our power simulations suggest that an X-QTL experiment employing DSPR RILs can be quite powerful if certain experimental parameters are carefully optimized.

### X-QTL mapping resolution can be high

Although the power of an X-QTL experiment can be high, if the approach suffers from poor QTL localization ability it may not be suitable for the dissection of complex traits. We used the actual genotypes of DSPR RILs (King *et al*. 2012b) to simulate a base population to be used for X-QTL mapping. Figure 3 depicts the average LOD profiles for a simulated QTL on chromosome 3L as a function of simulated parameters for a base population instantaneously expanded to a large size, randomly mated for four generations, and then phenotyped/genotyped in an eight-fold replicated experiment. Additional generations of random mating were also simulated and are discussed below. Localization is not greatly impacted by the number of individuals used in the DNA prep (*N_DNA_*), the number of RILs used to generate the base population (*N_RIL_*), or the selection intensity (*i*). As was clear above, power is considerably higher - and the LOD score at the QTL peak is greater - for larger values of *N_DNA_* and *i*, so these parameters should be maximized regardless. Figure S2A depicts the LOD profiles in a manner similar to Figure 3, except shows only a single realization of our simulation. This illustrates the considerable variation in the LOD profile of any particular simulation realization, or indeed of any real QTL mapping experiment.

**Figure 3:**
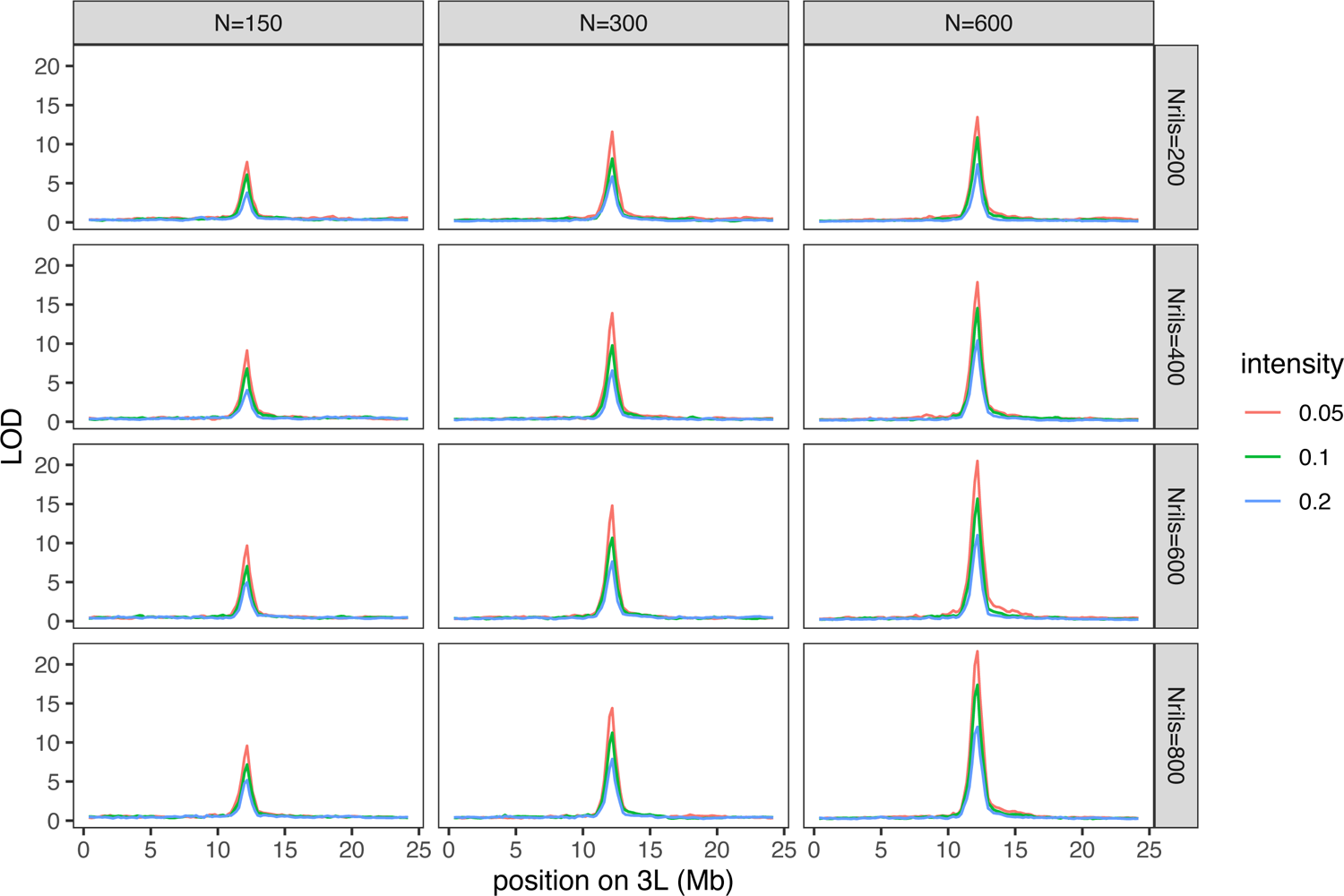
Average QTL localization for an X-QTL experiment. Simulated base populations were derived from DSPR population A RILs, and are created via diallel crosses of the RILs, instantaneous expansion, and then 4 generations of random mating. We only consider 200 positions spanning chromosome 3L, simulating a QTL contributing 2% heritability in the middle of the chromosome, and an eight-fold replicated experiment. Localization is presented as a function of the effective number of individuals contributing to the DNA pools (*N_DNA_* = 150, 300, 600), the number of RILs used to derive the base population (*N_RIL_* = 200, 400, 600, 800), and the selection intensity (*i* = 5%, 10%, 20%).

We carried out additional localization simulations of a smaller region, at a higher marker density, considering only DNA preps of 300 or 600 flies, and four generations of random mating following base population creation (focusing on a narrow set of parameters reduced computational effort considerably). Figure S2B plots the average 2-LOD support window size as a function of average QTL peak LOD score, with each point the average over 250 pure replicate simulations for any particular parameter combination. This relationship shows that localization ability is a strong function of peak LOD score, with highly significant peaks being better localized, much like QTL mapping in general. An ANOVA (see the GitHub site) examining the effect of different experimental parameters (*N_RIL_*, *N_DNA_*, *i,* and *N_rep_*) on localization after including average LOD score as a covariate, shows that only increasing the number of experimental replicates further impacts localization ability. That is, although the manipulated parameters all impact localization ability, their impact on localization largely parallels their impact on power. Figure 3 and Figure S2B demonstrate that in situations where the X-QTL design is powerful, there is considerable localization ability, in many cases approaching sub-cM QTL intervals. For more modestly-sized experiments localization ability is on the order of 1-cM, comparable to RIL-based mapping with >500 RILs and multiple individuals measured per RIL (King *et al*. 2012a).

### The impact of the number of founding RILs

It is not surprising that increasing the number of replicates of an experiment yields an increased LOD score at the QTL. It is also expected that increasing the number of flies contributing to the DNA pool similarly increases the average QTL LOD score; As the number of flies contributing to a DNA pool increases, haplotype frequency estimates in the pool approach those of the population from which the pool is sampled, thus increasing the power of the hypothesis test. It is also intuitive that increasing the selection intensity while holding *N_DNA_* constant will result in greater power. However, we further observe that the number of RILs used to create the base population has a large impact on power, particularly between 200 and 600 founding RILs (Figure 3); It is unclear why this is, since we might expect the number of founder RILs to primarily ensure founder alleles are represented in proportion to their frequency in the base population.

In our initial power simulations the false positive rate and significance threshold is obtained from a comparison of two equally-sized draws without phenotypic selection from the base population, with other parameters varied. In order to rule out the possibility that certain parameter combinations give only the appearance of increased power, we further examined the distribution of LOD scores at markers loosely linked to the causative site (*i.e.*, markers >7Mb from the causative site; see Figure S2C). Given the mapping resolution of our experiment, the distribution of LOD scores at positions distant from the QTL location should approach that of two equally-sized draws from the base population. However, LOD scores could be elevated at loosely-linked markers if a limited sample of founder RILs led to linkage disequilibrium between causative loci and the polygenic background.

In Figure S2C we observe slightly elevated LOD scores for certain high power parameter combinations (*i.e*., those employing strong selection, large pool sizes, and many replicates) when the number of generations of random intercrossing following base population establishment is <= 4, consistent with some X-QTL signal coming from polygenic background variation. Despite our simulations being able to detect this potential source of false positive mapping signal at loosely-linked sites, this factor appears to have only a subtle impact on the genomewide false positive rate, and the force is extremely weak for many realistic parameter combinations. It is also apparent that 4-8 generations of free recombination after establishment of the base population is very effective at removing this source of variation. More than 8 generations of free recombination results in little additional gain, and we speculate this is because recombination events in an expanding base population remain at low frequency, and do not have nearly the impact on localization as events present in the RILs.

Theoretical arguments aside, in practice a few generations of population expansion and recombination are likely unavoidable given the size of the base population necessary to select harshly (5% or 10% of individuals) while maintaining a large number of individuals in the selected pool (ideally up to 600). A corollary is that maintaining very large populations for extended periods is challenging, and can allow for significant haplotype frequency shifts (King et al. 2012), so there will likely be a tradeoff between ensuring several of generations of intercrossing to limit long-range LD, and avoiding major changes to haplotype frequencies via selection and/or drift.

### Accurate haplotype frequency estimation with modest genomewide sequencing coverage

Haplotype frequency estimates from pooled re-sequencing data with known founders can be quite accurate, depending on the window size, the number of founder haplotypes, the sequence divergence among haplotypes, and the level of recombination the population has experienced (Kessner *et al*. 2013; Tilk *et al*. 2019; Linder *et al*. 2020). We developed a method to estimate haplotype frequencies from pooled samples composed of known founders in the context of 4- and 18-way multiparental yeast populations (Cubillos *et al*. 2013; Burke *et al*. 2014b; Linder *et al*. 2020). In contrast to the yeast system, where the DNA pool consists of billions of cells/individuals, sampling in *Drosophila* is finite, and technical factors (body size differences, variation in the level of lysis in a bulk DNA isolation, etc) will further lower the effective population size in a pool of individuals. On the other hand, the SNP density relative to the size of non-recombined haplotype blocks can be higher in flies than in yeast. Furthermore, different yeast strains can share long, highly-similar genomic segments (Peter *et al*. 2018) making unique haplotype assignment difficult. As a result the problem is slightly different, with flies having both advantages and disadvantages relative to yeast. We sought to quantify the accuracy of haplotype frequency estimation in flies for the particular founders and RILs actually employed.

Although we cannot measure the effective pool size in any *Drosophila* X-QTL experiment, we can measure the average squared difference between two estimates of each founder’s haplotype frequency; the frequency of SNPs private to a single founder in a high coverage (240X) dataset, and the haplotype frequency estimate for the window nearest this SNP from down-sampled data. We detail this experiment in the Materials & Methods to maintain readability. The principal result is that the average squared difference over millions of private SNPs is not very sensitive to the level of downsampling, with estimates of haplotype frequency being extremely accurate at ∼35X coverage. Furthermore, based on the variance in SNP frequency estimates due to sampling, we estimate that the average absolute error in allele frequency from the haplotype caller is ∼1%. This error is approximately constant for coverages from 17X to 240X.

It is important to note, and is perhaps counter-intuitive, that under binomial sampling one would need to sequence to ∼1000X average coverage to estimate SNP frequencies with the degree of accuracy we achieve for haplotypes with only 35X coverage. It is also important to note that the errors on haplotype frequency estimates are small enough (even at 35X coverage) that the finite number of flies contributing to a DNA pool can contribute to noise and lack of power. That is, with 500 individuals equally contributing DNA to a pool, the sampling error on the estimate of true population frequencies are the same order of magnitude as the error in haplotype estimation, supporting our claim that it is important to carry out X-QTL experiments using large pools of 500-1000 flies. Counter-intuitively the sequencing of pools can likely be carried out at modest and cost-effective coverage with only subtle effects on the outcome.

### Experimental test of X-QTL mapping identifies multiple QTL

We carried out an X-QTL experiment using a population constructed by mixing 663 DSPR pA RILs, allowing the population to undergo free recombination for 1-5 generations, and selecting for caffeine resistant females in a 4-fold replicated experiment. Each replicate tested 2,337-2,572 animals, selecting the 9.0-10.3% most caffeine resistant females in each case (228-254 animals), along with a sample of 250 control animals from the base population. Full details of each replicate are presented in Table S1. We extracted DNA in bulk from each of the 8 resulting samples, constructed libraries, and generated ∼140X of alignable sequence per pool. We estimated frequencies of the 8 haplotypes in each pool over 200-kb windows with a 10-kb step size, and for each window carried out a linear model testing for a change in the arcsin square root allele frequency between the selected and control pools (see Materials & Methods for full details).

Unlike the case of RIL- or individual-based QTL mapping, it is unclear how to create a permutation-based null distribution for X-QTL mapping (Doerge and Churchill 1996). We carried out simulations comparing haplotype frequencies in equally-sized draws from control populations and examined the distribution of LOD scores as a function of experimental parameters. Over all parameter combinations tested, a LOD threshold of 4 held the genomewide false positive rate below 5%, and is a reasonable threshold to employ. This being said, this threshold may be slightly anti-conservative if the number of flies contributing to the DNA pool is less than 500, as was the case for our caffeine resistance experiment. To further investigate an appropriate threshold we simulated a dataset that mimicked the precise caffeine experiment carried out (effective number of individuals in the DNA pool = 150, number of replicates = 4, selection intensity = 10%, no QTL) and created QQ plots (Figure S3). Pools of ∼250 female flies (effective *N_DNA_* = 150) can be associated with a slightly elevated false positive rate, and this is observed in the QQ plot with the simulated data showing some inflation. Nonetheless, for this parameter combination a LOD threshold of 4 is associated with an acceptable genomewide false positive rate of 0.22%. Figure S3 also plots the distribution of observed LOD scores, and LOD scores after removal of all test statistics within 2-Mb of each of our 7 significant QTL peaks (Table 1 - discussed below). The LOD scores associated with actual scans - with or without peak regions removed - show much greater inflation. The simplest explanation for inflation after mapped QTL regions are removed is that the signal of linkage extends over regions larger than 2-Mb, and that QTL of subtle effect that do not reach the threshold for significance are contributing to signal throughout the genome.

Figure 4 plots the genomewide −log_10_(*P*) values comparing haplotype frequencies in caffeine-selected and control pools based on the full-coverage (∼142X) pooled sequencing dataset, as well as for datasets downsampled to ∼36X or ∼14X. Two patterns are immediately apparent: First, we see several significant, well-localized peaks (Table 1), consistent with our power and localization simulations (Figures 2 and 3). It appears we observe significant QTL despite our experiment being non-optimal in terms of the number of replicates, the selection intensity, and the number of flies in each DNA pool. These differences were due to our analytical and experimental approaches both moving forward contemporaneously, and because we elected to employ a fairly challenging caffeine resistance phenotyping regime for X-QTL mapping to more closely mimic that used originally in RILs (Najarro *et al*. 2015); Loading individual flies into single *Drosophila* activity monitor tubes (as opposed to bulk testing multiple individuals in standard fly vials) was only done to copy our previous work. The benefit of our choice of character is that we are in a position to directly compare mapped QTL identified via a pooled population approach and a RIL-based approach.

**Figure 4:**
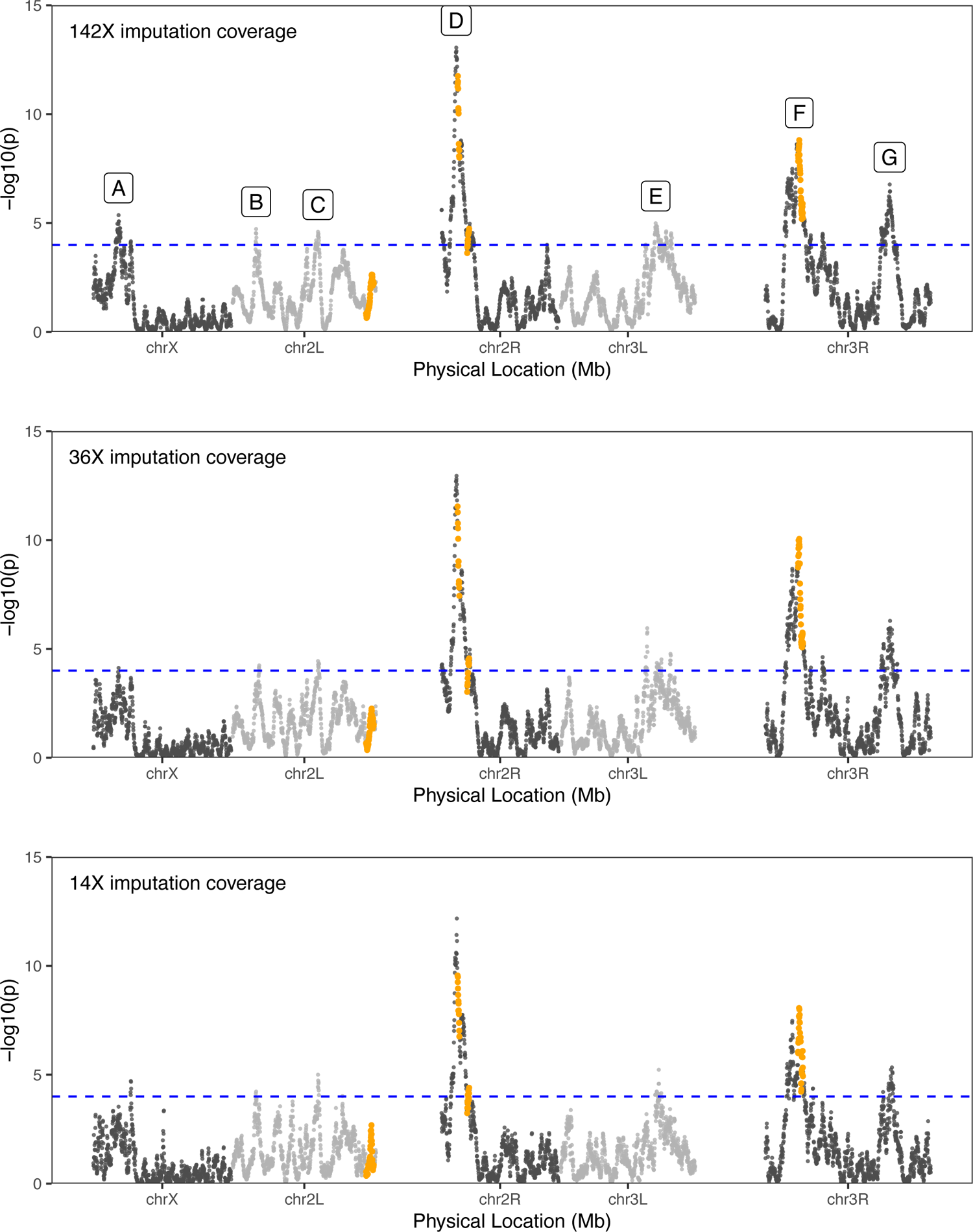
X-QTL scan for caffeine resistance loci. Each panel shows −log_10_(*P*) test statistics derived from comparisons between caffeine-selected and control pools using the full coverage pooled sequencing data (∼142X per sample), and data where read coverage was downsampled to ∼36X or ∼14X. Points on different chromosome arms are shaded differently, the x-axis scale is proportional to physical location using the *D. melanogaster* reference Release 6 genome coordinates, and heterochromatic regions (Table S3) are not plotted (leading to gaps at centromeres). The statistical threshold for QTL detection (horizontal blue, dashed line) is set at −log_10_(*P*) = 4. Each called QTL “peak” is given a letter code, corresponding to codes in Table 1 and Figure 5. Orange points correspond to the 2-LOD drop intervals of caffeine resistance QTL previously mapped by Najarro et al. (Najarro *et al*. 2015) using the DSPR pA RILs directly.

Second, the locations and significance levels associated with each peak are very similar even for highly downsampled read data (compare the 3 panels of Figure 4). This is consistent with our earlier result that error in haplotype frequency estimates is largely independent of coverage over the explored range. That the downsampled datasets yield similar results to the full dataset suggests future experiments can be more efficiently and inexpensively carried out at 20-40X sequencing per pool. This may be counterintuitive to some, but is consistent with other published claims that when founders are known, haplotype frequency estimation from pooled sequencing data can be quite accurate even with low coverage read data (Kessner *et al*. 2013; Tilk *et al*. 2019; Linder *et al*. 2020).

Figure 5 presents LOD scores and allele frequency changes at QTL, and Table 1 provides the properties of these mapped QTL, for the full coverage (∼142X) dataset. Figure S5 presents mapped QTL and frequency changes using the ∼35X dataset. It is apparent that the properties of mapped QTL are largely the same, except for X-QTL:E which shifts roughly 1-Mb to the left when using the downsampled data; the X-QTL:E region has two “peaks” that only just reach significance depending on the pooled sequencing dataset employed, and has a fairly wide 2-LOD support interval. The *overall* genomewide similarity of the LOD profiles, especially for the more significant regions, supports our claim that little (but some) additional information is gained by sequencing libraries to higher coverage. If this experiment were repeated the impact of additional experimental replicates, stronger selection, or more flies in the selected pools would likely dwarf that of additional sequence coverage (see above simulation results).

**Figure 5:**
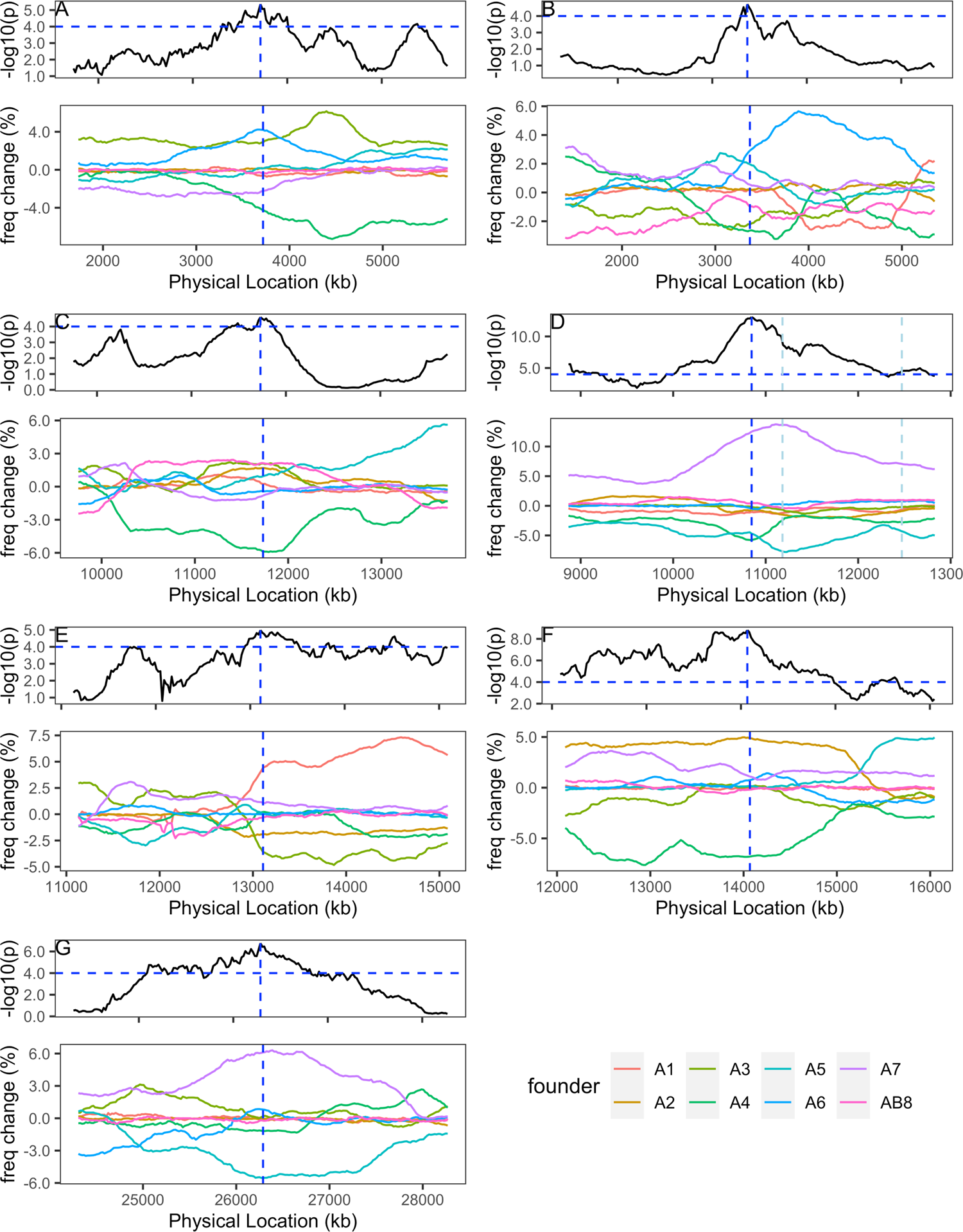
Founder haplotype frequencies at caffeine X-QTL. A 2-Mb window of the genome, centered on each of the X-QTL loci mapped (A-F, see Figure 4 and Table 1) is presented for the full coverage sequencing data. The top panel for each X-QTL presents the −log_10_(*P*) test statistics across the window, with the vertical dashed blue line at the QTL peak, and the horizontal dashed blue line at LOD=4. The bottom panel for each X-QTL presents the change in frequency of each haplotype, polarized such that positive values correspond to an increase in the frequency of that haplotype in the selected pool. In panel X-QTL:D the vertical light blue lines represent the locations of the Q2 (left) and Q3 (right) loci mapped by Najarro et al. (Najarro et al. 2015) illustrating the physical proximity of these loci, and the similarity of the haplotype frequencies, which together suggest that Q3 of the prior work may be spurious.

### X-QTL replicates a subset of mapped caffeine resistance loci

Najarro et al. (Najarro *et al*. 2015) resolved several caffeine resistance loci via directly screening DSPR RILs, four of which were mapped in the pA population: Q1 mapped to chromosome 2L adjacent to the centromere (LOD = 9.9, percent heritability explained = 5.5%), Q2 mapped to the centromeric end of 2R (LOD = 27.2, heritability = 14.4%), Q3 mapped ∼2-cM to the right of Q2 on 2R (LOD = 11.3, heritability = 6.3%), and Q9 mapped to 3R (LOD = 10.2, heritability = 5.7%). The 2-LOD drop intervals of these 4 QTL are reflected in our X-QTL map with orange symbols, and it appears that we replicate 2 of them (Figure 4, top panel, X-QTL:D and X-QTL:F).

The largest-effect QTL mapped in RILs, Q2, was replicated by X-QTL:D (Figure 4, Table 1). Najarro et al. (Najarro *et al*. 2015) found that RILs carrying the A7 haplotype at this locus had the highest average caffeine resistance. Similarly, we estimate that at this position in our X-QTL study haplotype A7 exhibits the greatest frequency increase in the selected pools over the control pools (Figure 5D). Notably, founder A7 is the only pA founder strain that possesses 2 copies of the *Cyp12d1* cytochrome P450 detoxification gene (as does the *D. melanogaster* genome reference strain, where the copies are termed *Cyp12d1-d* and *Cyp12d1-p*). *Cyp12d1* expression is known to be induced in response to caffeine (Willoughby *et al*. 2006; Coelho *et al*. 2015; Najarro *et al*. 2015), and experiments using ubiquitous Gal4-UAS-RNAi knockdown (under the control of an actin promoter), and adult-specific knockdown (using an RU486-inducible “GeneSwitch” actin Gal4 driver) indicated that *Cyp12d1* knockdown reduces caffeine resistance (Najarro et al. 2015). Additionally, Najarro (Najarro *et al*. 2015) found a significant association between *Cyp12d1* copy number variation (CNV) and caffeine resistance in both the pA and pB RIL panels. By controlling for *Cyp12d1* CNV status during RIL-based QTL mapping (via the addition of a covariate into the mapping model) the Q2 QTL was eliminated, suggesting the CNV or something in linkage disequilibrium with it - is causative.

We also replicated RIL-based QTL Q9 with X-QTL:F (Figure 4, Table 1). In our previous RIL-based study, RILs carrying founder haplotype A2 at this location showed higher average caffeine resistance, while RILs carrying A4 had among the lowest resistance. We recapitulate these findings, showing that A2 exhibits the greatest frequency increase in selected pools, while A4 shows a large frequency decrease in the selected pools (Figure 5F). Both the Q9 and X-QTL:F intervals contain *Cyp6d5*, a gene that is transcriptionally induced in response to caffeine exposure (Najarro *et al*. 2015).

Q1 from our RIL-based study does not replicate here. We previously estimated its effect to be modest, so easy replication may not be anticipated (Zhou *et al*. 2020). Additionally, the QTL is extremely close to the chromosome 2 centromere, a location where effective mapping is challenging (Noor *et al*. 2001); The original Q1 may not represent a true causative caffeine resistance locus. RIL-based Q3 also fails to replicate, and again it has a modest effect. In addition, Najarro et al. (Najarro *et al*. 2015) mapped Q3 to a position just 2.5-cM away from the large-effect Q2 QTL, so Q3 may not represent an independent locus; Indeed the haplotype frequencies at both these positions are similar in our X-QTL populations (Figure 5D, compare frequencies at positions 11,182-kb and 12,472-kb, which are the locations of Q2 and Q3, respectively).

### Functional testing plausible candidate caffeine resistance genes

Six of the 7 X-QTL we identify are mapped to 440-920kb intervals, each encompassing 41-105 protein-coding genes (Table 1). In addition to the two candidates discussed above - *Cyp12d1* (X-QTL:D) and *Cyp6d5* (X-QTL:F) - a number of plausible, novel candidate genes emerge from mapped X-QTL. X-QTL:A includes *Tlk* (*Tousled-like kinase*), a gene involved in cell cycle progression and the DNA damage response, that also contains a caffeine-sensitive phosphorylation site (Groth *et al*. 2003). X-QTL:B includes *E23* (*Early gene at 23*), which encodes an ATP-binding cassette (ABC) transporter subunit (Hock *et al*. 2000) - ABC transporters have important roles in xenobiotic metabolism (Xu *et al*. 2005), and *Ugt36A1* (*UDP-glycosyltransferase family 36 member A1*) which is a UDP glycosyltransferase detoxification enzyme gene. X-QTL:C encompasses *Crys* (*Crystallin*), whose gene product is involved in the formation of the peritrophic matrix (Kuraishi *et al*. 2011), a structure that forms a protective layer lining the midgut epithelium (Terra 2001), and *Vha100-5* (*Vacuolar H^+^ ATPase 100kD subunit 5*) which encodes a subunit of an ATP-dependent proton pump, and is expressed in the fly midgut (Overend *et al*. 2016). X-QTL:G encompasses the gene *osy* (*oskyddad*) which encodes an ABC transporter subunit, is required to provide a barrier to xenobiotics in *D. melanogaster* larvae (Wang *et al*. 2020), and shows reduced gene expression in adult males exposed to caffeine (Coelho *et al*. 2015).

To evaluate the effects of each of these 8 genes we employed both ubiquitous RNAi knockdown using Gal4 under the control of an actin promoter, and expressed Gal4 specifically in the anterior portion of the adult midgut. Using both drivers, and both reciprocal cross directions (see Materials & Methods), both *Cyp12d1* and *Cyp6d5* show a significant reduction in lifespan on caffeine-containing media compared to the control, no-knockdown genotype (Dunnett’s tests, 0.05 < *P* < 10^−12^, Figure 6), re-confirming the strong relevance of this pair of detoxification genes in the response to caffeine challenge. The *Tlk* gene (X-QTL:A) only shows a significant reduction in resistance when knocked down in the gut in 1 of the 2 reciprocal crosses (cross direction-specific Dunnett’s tests, *P* < 10^−4^ and *P* = 0.08), while the ubiquitous knockdown was lethal (Table S2). Of the two genes tested within the X-QTL:B interval (*E23*, *Ugt36A1*) the *E23* ABC transporter subunit gene appears to be the best candidate based on the RNAi; Knockdown of this gene leads to significantly reduced lifespan in all 4 experiments (Dunnett’s tests, 0.01 < *P* < 10^−8^, Figure 6), while knockdown of *Ugt36A1* shows no effect on phenotype in the gut, and only a minor reduction in resistance in 1 of the 2 ubiquitous knockdown reciprocal crosses. Of the two genes tested beneath X-QTL:C, *Crys* shows no evidence for an effect (Figure 6), while *Vha100-5* gut knockdown reduces resistance (cross direction-specific Dunnett’s tests, *P* < 10^−4^ and *P* < 0.01), and resistance is reduced in 1 of the 2 reciprocal crosses of the ubiquitous knockdown tests (Dunnett’s tests, *P* = 0.8 and *P* < 0.001). Finally, *osy*, the sole gene tested within the X-QTL:G interval shows no effect with gut knockdown (Figure 6), and ubiquitous knockdown led to lethality (Table S2). Ultimately, the novel caffeine resistance candidates *E23* and *Vha100-5* appear particularly worthy of additional functional study.

**Figure 6:**
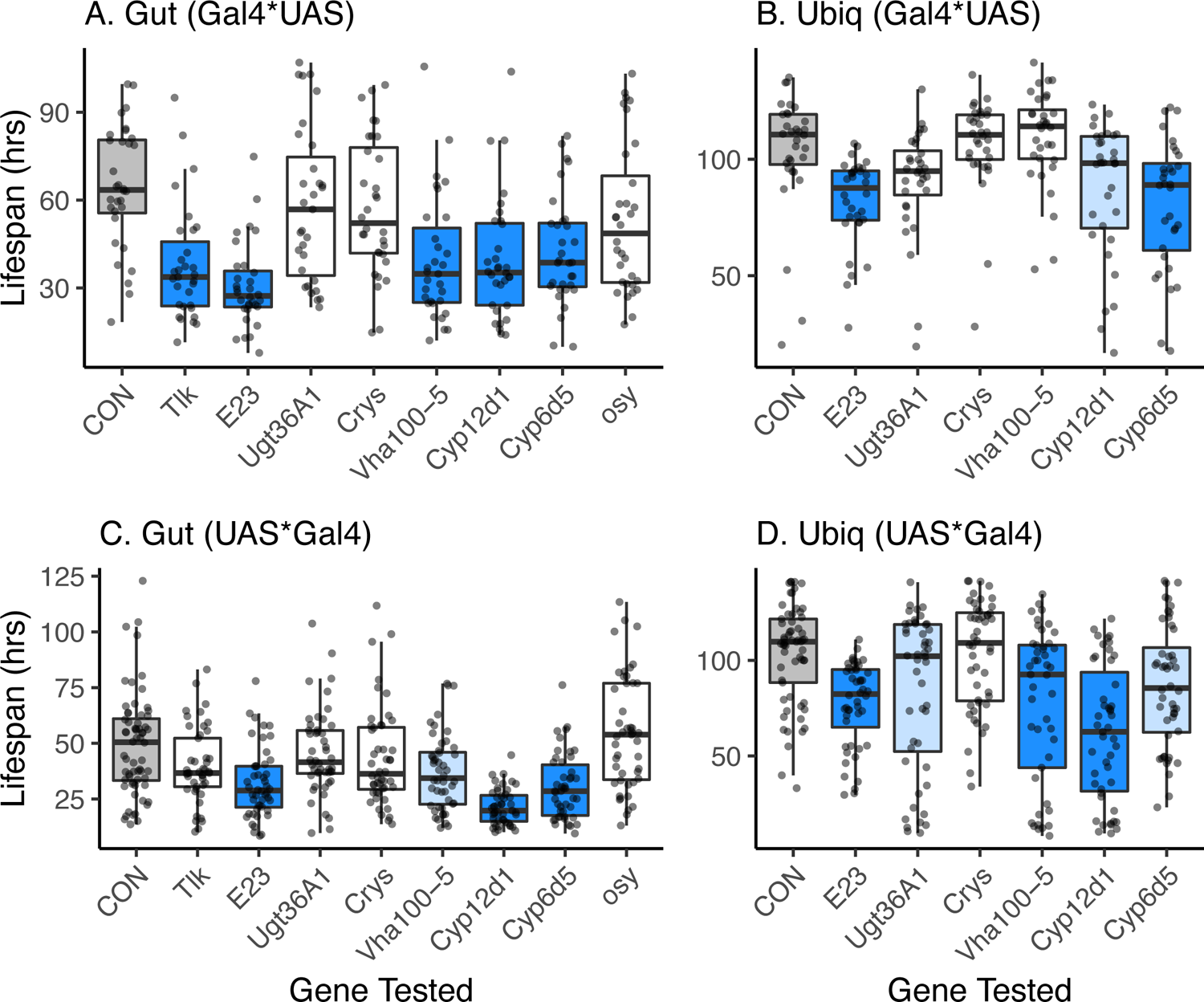
Effects of candidate gene RNAi knockdown on caffeine resistance. Shows the lifespan (in hours) of replicate females from a series of Gal4-UAS-RNAi genotypes (and controls, CON) on 1% caffeine media. The genes impacted are listed by their symbols along the x-axes of the plots. Each dot represents the phenotype of a single individual, and for each genotype the individual-level scores are overlaid with a boxplot (following the R ggplot2 ‘geom_boxplot’ defaults). Each panel represents a different Gal4 driver and a different cross direction: (A and C) anterior midgut driver, (B and D) ubiquitous driver. (A and B) Test individuals are the result of crossing male Gal4 to female UAS, (C and D) test individuals are the result of crossing male UAS to female Gal4. Within each combination of driver and cross direction, all Gal4-UAS-RNAi genotypes were compared to the control genotype using a Dunnett’s test. Box colors reflect these tests: gray = CON, white = not significantly different from control (*P* > 0.05), light blue = significantly different from control at *P* < 0.05, dark blue = significantly different from control at *P* < 0.005.

## DISCUSSION

Here we explore an alternative to RIL-based QTL mapping in *Drosophila* that is analogous to the extreme QTL (or X-QTL) approach originally conceived of in yeast (Ehrenreich *et al*. 2012), and more recently used in the nematode model system *C. elegans* (Burga *et al*. 2019). Conceptually derived from the bulked segregant analysis mapping strategy (Michelmore *et al*. 1991), X-QTL mapping as employed in yeast utilizes extremely large pools of haploid segregants from a single generation cross between two strains. Unlike this two-parental strain, single-generation cross, the base population we employ is created by combining several hundred advanced intercross DSPR RILs (King *et al*. 2012a; b), and up to eight haplotypes segregate at any given genomic location in the DSPR X-QTL mapping population.

X-QTL mapping draws replicate samples of phenotypically-extreme and control individuals from a large base population, and identifies regions of the genome exhibiting consistent differences in founder haplotype frequencies between extreme and control pools following short read sequencing (Figure 1). For a QTL contributing 2% to variation in a complex trait we explore the power and localization ability of this X-QTL mapping approach as a function of the number of experimental replicates, the number of RILs used to found the base population, the number of individuals contributing to a DNA pool, the selection intensity employed, and the number of generations of expansion/recombination prior to phenotyping the intercrossed base population. Our simulations suggest that for difficult, but experimentally achievable designs the X-QTL approach can routinely have considerably higher power than traditional RIL-based mapping (Figure 2), and at least as high QTL localization ability (Figure 3). This being said, power and localization ability is impacted by the precise experimental design employed and the highest power is only achieved when the number of RILs contributing to a base is greater than 500, selection intensity is greater than 10% (ideally one should aim for ∼5%), the number of flies contributing to the DNA pool is >500 (implying that the total number of individuals assayed is >=5000), and four (and ideally 8 or more) experimental replicates are used.

Several other mixed population mapping strategies have previously been executed in *Drosophila*. Huang et al. (Huang *et al*. 2012) derived a population - termed “Flyland” - from 40 inbred strains of the DGRP (Mackay *et al*. 2012; Huang *et al*. 2014). This population was allowed to undergo recombination for 70 generations, 2,000 individuals were phenotyped for each of three complex traits, and allele frequencies were contrasted between sequenced pools of 300 phenotypically-extreme individuals. A similar design has been employed in studies of a number of other phenotypes (e.g. (Morozova *et al*. 2015; Zhou *et al*. 2017; Fochler *et al*. 2017)), each resulting in tens to hundreds of variants exhibiting allele frequency differences between control and selected pools. No analytical details of the power or resolution of this approach are available, and in contrast to our simulated and experimental data, where increasing numbers of replicates improve power, these studies are generally un-replicated. Flyland studies also typically phenotype fewer individuals than our simulations suggest is optimal for the highest power. And since the strategy relies on directly contrasting SNP allele frequency between pools, rather than use the haplotype approach we take (which improves allele frequency estimates, (Tilk *et al*. 2019)), unless sequencing coverage is high, the power of the Flyland approach may be low.

Another group has proposed the use of “hybrid swarm” populations. Here, many inbred strains 34 in the case of Erickson et al. (Erickson *et al*. 2020) - are mixed for 4-5 generations, then recombinant animals are phenotyped and individually subjected to ultra-low pass sequencing, facilitating GWAS-type mapping analyses (Erickson *et al*. 2020; Weller *et al*. 2021). This approach has the advantage of not requiring RILs, and needing only a small number of generations of population maintenance. But, while the total sequencing effort is reasonable, rather than working with DNA pools, an investigator would need to efficiently and inexpensively generate thousands of individually-indexed sequencing libraries.

Several studies have used various mixed population designs to genetically dissect body pigmentation, a classic *Drosophila* trait impacted by a small set of well-documented candidate genes (reviewed in (Massey and Wittkopp 2016). Bastide et al (Bastide *et al*. 2013) phenotyped 8,000 outbred animals derived from thousands of unknown founders, and after sequencing pools of extremely light or dark animals, successfully resolved large effects to classic pigmentation loci (*bric-a-brac*, *ebony*, *tan*). Dembeck et al (Dembeck *et al*. 2015) created a population from 3 light and 3 dark pigmented DGRP lines, maintained this for a few generations, and found allele frequency differences between sequencing pools generated from light and dark flies that localized to classic pigmentation gene locations, in addition to other regions of the genome. Bastide et al. (Bastide *et al*. 2016) used a series of two-way advanced intercross, bulked segregant analysis mapping populations to yield a similar result; Some peaks resolve to intervals harboring known pigmentation genes, while others might implicate novel candidates. Collectively, these studies suggest there are intermediate frequency causative SNPs of large effect impacting *Drosophila* pigmentation, and as a result frequency differences at single SNPs in mapping populations can be dramatic, and the effects at QTL sometimes be quite large. The applicability of these approaches to more polygenic traits is unknown.

Perhaps the most similar experiment to the one we describe here is Burke et al (Burke *et al*. 2014a), who sequenced pools of the 2% most extreme long-lived individuals in the synthetic populations from which the DSPR RILs were derived (King *et al*. 2012b). However, this aging study only obtained ∼100 flies for each sequencing pool, and while 4 populations were used, none were replicated. The simulations we present here suggest that despite some success, this longevity experiment was modestly powered.

Key features of the X-QTL approach we recommend are that a large, highly-recombinant population is created, and that several pairs of matched extreme and control DNA pools are each generated from large numbers of animals, and sequenced. A great deal of the power of the design comes from the populations being derived from a modest number of known founders, which allows tests for differentiation between control and extreme pools to be carried out on imputed founder haplotype frequencies, as opposed to using directly ascertained SNP frequencies.

A perhaps non-intuitive, but well-established result (Long *et al*. 2011; Kessner *et al*. 2013; Burke *et al*. 2014b; Tilk *et al*. 2019; Linder *et al*. 2020) is that by virtue of the base population ultimately consisting of genetic material from only 8 highly characterized “founders”, sliding window haplotype frequency estimates are extremely accurate, even with as little as 20-40X of short read sequencing. We show above that at 35X sequencing coverage, for much of the genome, founder haplotype frequencies are estimated with errors of ∼0.01. Accurate haplotype frequency estimates strongly impact the power of the X-QTL approach in flies. If founder haplotypes are ignored or unavailable, and SNP frequencies are employed directly, one would require sequencing coverage of ∼1000X to obtain similarly accurate allele frequency estimates. The modest sequencing coverage necessary under our design means that the sequencing budget for an 8-fold replicated experiment can be relatively inexpensive. A primary practical limitation of our approach is fly handling; the pools of flies used to make DNA must consist of 100’s of individuals, and with the necessary harsh selection regime, thousands of individuals must initially be targeted.

Our 4-fold replicated proof of principle experiment used pools of ∼250 female flies either randomly chosen from a base population, or in the top ∼10% of individuals surviving following exposure to high levels of caffeine. We chose this demonstration trait as it has been the subject of past research using a RIL-based mapping approach (Najarro *et al*. 2015), and we wished to determine if QTL identified by the two methods mapped to the same locations. This being said, the assay method involves considerable fly handling and is not ideally suited to X-QTL mapping. As a result, our empirical test of the approach was carried out with parameters near the lower end of those our simulations suggest yield the highest power. Nonetheless, we still mapped 7 X-QTL at a genomewide false positive rate of <5% (Table 1). Two of these X-QTL overlap with, and appear to replicate 2 of the 4 QTL mapped by Najarro et al. (Najarro *et al*. 2015) in the population A panel of the DSPR RILs, the progenitors of the X-QTL population employed in the present study. Additionally, the X-QTL results suggest that one or both of the previously-mapped, but un-replicated QTL may be spurious; one is very close to the centromere, and the other is very close to a larger-effect, replicated locus. Notably, we were able to map 5 new QTL associated with 6 novel and pre-existing candidate genes (Table 1). Using both general and gut-specific Gal4 drivers, we attempted to functionally validate the implicated genes; Both candidates -*Cyp12d1* and *Cyp6d5* -underlying the pair of replicated QTL found here and in Najarro et al. (Najarro *et al*. 2015) yielded reduced caffeine resistance on knockdown. Furthermore, two new candidate genes -*E23* and *Vha100-5* -received functional support.

Comparing two caffeine resistance experiments employing near identical phenotyping regimes, it appears the results from the X-QTL study provided at least similar, if not superior genetic insight into the phenotype than the RIL-based study. And had we not wished to compare designs, enforcing use of the same phenotyping assay, a less cumbersome bulked phenotyping approach would likely have been possible for X-QTL mapping (e.g. exposing groups of flies to caffeine-supplemented media in vials versus testing flies singly in narrow tubes), yielding larger numbers of phenotyped animals, and -as our simulations suggest -higher power.

Several caveats of X-QTL mapping in flies are apparent, many resulting from the pooled nature of the phenotyping and genotyping that is employed. First, since X-QTL are detected as haplotype frequency differences between pooled samples each consisting of hundreds of outbred animals, one does not obtain estimates of dominance or epistasis associated with mapped X-QTL. Dominance and epistasis can be estimated using RILs (or individuals), although we note dominance cannot be estimated when RILs are directly phenotyped, the most common mapping strategy using RILs. Furthermore, in MPPs segregating for 8 founder alleles, allele-specific dominance and epistatic terms can be difficult to accurately estimate. Working directly with 8-way RILs there are 64 two gene epistatic terms, and if the RILs are intercrossed to obtain heterozygous genotypes there are potentially 28 dominance terms, and a very large number of epistatic terms. Investigators typically measure on the order of ∼500 DSPR RILs, so estimating dominance and epistasis in the MPP framework can quickly result in over-fitted models. The result is that in practice, dominance and epistasis are rarely studied using the DSPR (although see (King *et al*. 2012b; Cogni *et al*. 2016)). Finally, while any characterization of the genetic basis of complex trait variation will be incomplete without a description of dominance and epistasis (Ehrenreich 2017), the contribution of non-additive effects to the variance of complex traits may not be high in most cases (Schizophrenia Working Group of the Psychiatric Genomics Consortium 2014; Bloom *et al*. 2015; Albert *et al*. 2018; Nag *et al*. 2020; Hivert *et al*. 2021). Further, when epistatic interactions have been identified in powerful yeast QTL studies, they generally involve at least one locus that also has a main effect (Bloom *et al*. 2015; Albert *et al*. 2018), implying that an X-QTL strategy can identify major players involved in epistatic interactions, and follow-up studies could identify epistasis.

Second, in a study using inbred lines one might collect multiple phenotypes on the same set of lines, allowing examination of genetic correlations among traits (e.g., (Dickson *et al*. 2016; Everman *et al*. 2019)). Furthermore, with inbred lines it is straightforward to include known covariates associated with each strain, for instance, infection status with the common *Wolbachia* microbe, or inversion genotype, during mapping (both these covariates are accounted for as standard by the DGRP analytical machinery; (Huang *et al*. 2014)). Equally, one can statistically account for larger-effect QTL, enabling more powerful mapping of smaller-effect loci (e.g., (Cogni *et al*. 2016)). Such analyses are not possible with X-QTL mapping given the pooled nature of the phenotyping and genotyping.

Third, screens of the mouse Collaborative Cross set of 8-way RILs have identified single RILs having particularly extreme or interesting phenotypes. For example, RILs having high susceptibility to epileptic seizures (Gu *et al*. 2020), or high susceptibility to *Salmonella* infection (Zhang *et al*. 2018). Such strains have potential utility as novel models of disease. But again, given the bulked phenotyping and genotyping approach that X-QTL relies on, such interesting phenotypes would not be captured in stable genotypes via X-QTL mapping.

Finally, it is clear that not all target traits will be as amenable to bulk phenotyping as the resistance trait we empirically explore here; when every individual must be individually handled/scored (e.g. counting bristle number - (Macdonald and Long 2007)) a RIL-by-RIL approach may be more profitable, especially given some of the other caveats of X-QTL mapping (above). However, X-QTL mapping is attractive, and can have high detection and localization ability for traits where some form of “self-selection” is possible (e.g., measures of toxin/stress resistance, and behaviors such as negative geotaxis, or oviposition preference). The list of traits that have been dissected using the DGRP (Table 3 of (Mackay and Huang 2018)) suggest that roughly half of the studied traits could have been phenotyped in bulk, some perhaps quite easily. That is, many of the traits Drosophilists currently study are potentially suitable for dissection via an X-QTL approach.

Despite some negatives, several additional efficiencies are obtained with X-QTL mapping. By virtue of working with a single large population, as opposed to several thousand vials, the accurate tracking of individuals/vials/blocks/strains necessitated in a RIL-based study is largely avoided. Furthermore, since phenotypes are all obtained from outbred genotypes, the impact of rare recessive genotypes and inbreeding depression are muted. In this regard, the contribution of inbreeding depression to RIL-based mapping results has rarely been quantified. Furthermore, as all flies are reared in a common garden, X-QTL mapping can also be advantageous for traits for which block- and/or vial-effects are unavoidable. Lastly, our simulations suggest that larger experiments can enjoy near single-gene mapping resolution, potentially dispensing with cM-sized confidence intervals and sorting through dozens of candidate genes.

It is important to note that the resolution of the experiments we simulate and experimentally employ assume the base population is initiated with highly-recombinant genotypes. DSPR RILs were derived from a population allowed to intercross for 50 generations prior to RIL initiation, and less recombined material would be associated with much lower mapping resolution. That said, the availability of the DSPR need not be a constraint on the design we outline. Indeed, early MPP-based mapping in *D. melanogaster* did not employ RILs, and instead directly interrogated a segregating population (Macdonald and Long 2007), much like the mouse Diversity Outbred population (Svenson *et al*. 2012). It would be possible to develop equivalent mixed base populations for X-QTL mapping by intercrossing the original 8 DSPR founder strains (or any set of inbred strains), followed by maintenance of the resulting population for 50 generations (∼2 years) to build up recombination events. Clearly the availability of RILs, and the ability to skip this population creation step, is a time- and labor-saving feature of the approach we outline.

X-QTL mapping can provide powerful, high-resolution mapping of QTL. In concert with emerging CRISPR-Cas9 homologous recombination-based allele replacements (Gratz *et al*. 2013, 2014; Ren *et al*. 2013; Port *et al*. 2014; Lamb *et al*. 2017), prime-editing strategies (Anzalone *et al*. 2019; Bosch *et al*. 2021), and recombinase-mediated cassette exchange following Cas9 (Bateman *et al*. 2006; Voutev and Mann 2018), X-QTL mapping may allow the field to move from mapped, main effect QTL to a more precise functional characterization of candidate genes than is possible via RNAi knockdowns. As these replacement/editing technologies become more mature, especially in model systems, QTL mapping may become more focused on the identification of high-confidence candidate genes underlying large, main-effect loci, with the accurate estimation of the effects associated with mapped factors being left to specific follow-up experiments. For many characters of interest to *Drosophila* geneticists the X-QTL approach we describe may provide a blueprint for quickly and cost-effectively identifying candidate genes underlying additive QTL.

## Acknowledgements

This work was supported by NIH R01 OD010974 (to SJM and ADL), NIH R01 ES029922 (to SJM), and NIH R01 GM115562 (to ADL). We also thank the University of Kansas Genome Sequencing Core facility (funded by NIH P20 GM103638) for library construction, and the Kansas INBRE program (funded via NIH P20 GM103418) for computational support.

## Figures Legends (Supplement)

**Figure S1:**
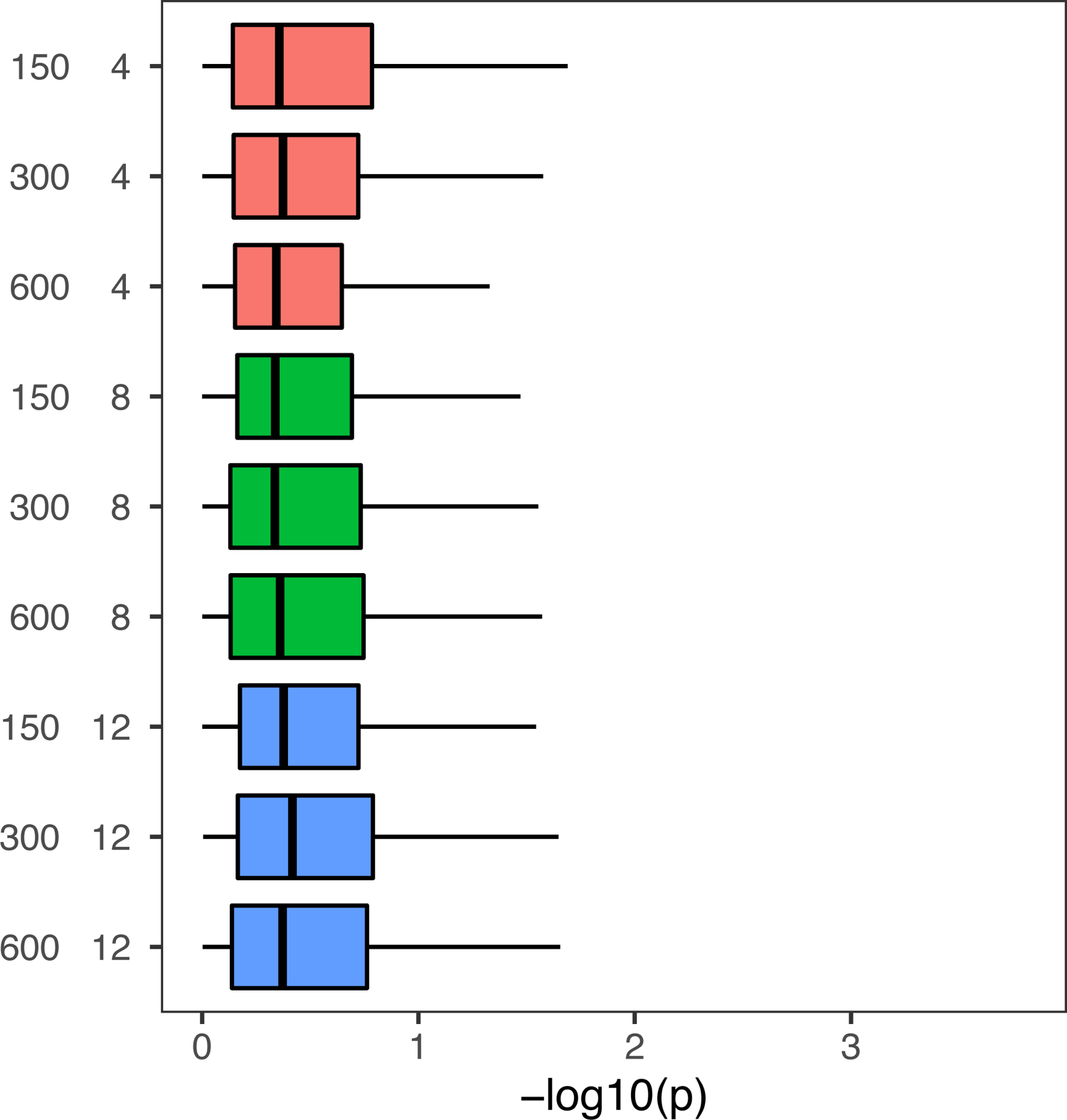
Null distribution of −log_10_(*P*) test statistics with X-QTL mapping. The expected distribution of the test statistic when comparing haplotype frequencies between two, equally-sized draws (of 150, 300, or 600 individuals) from the base population for each replicate (4, 8, or 12).

**Figure S2:**
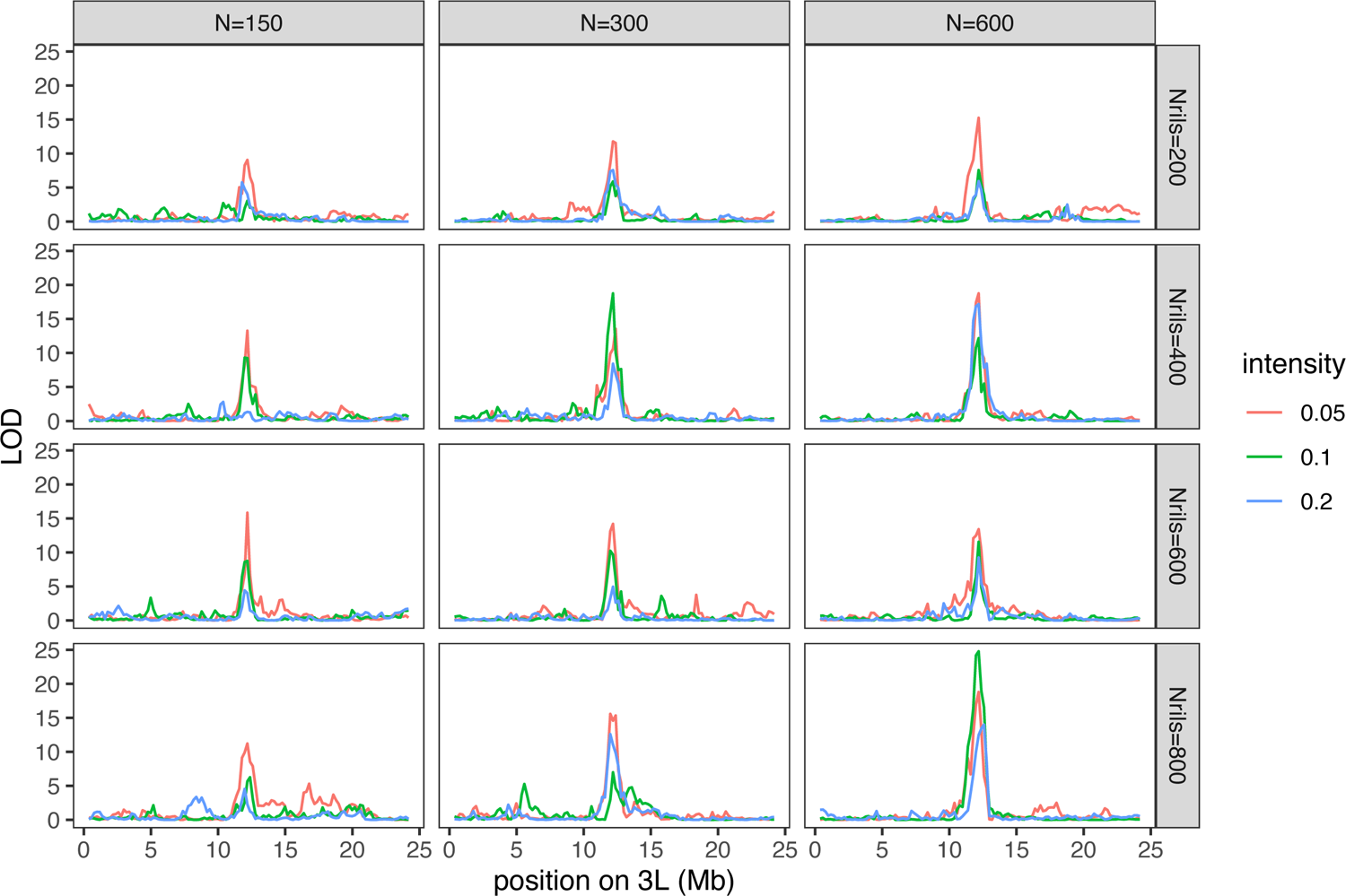
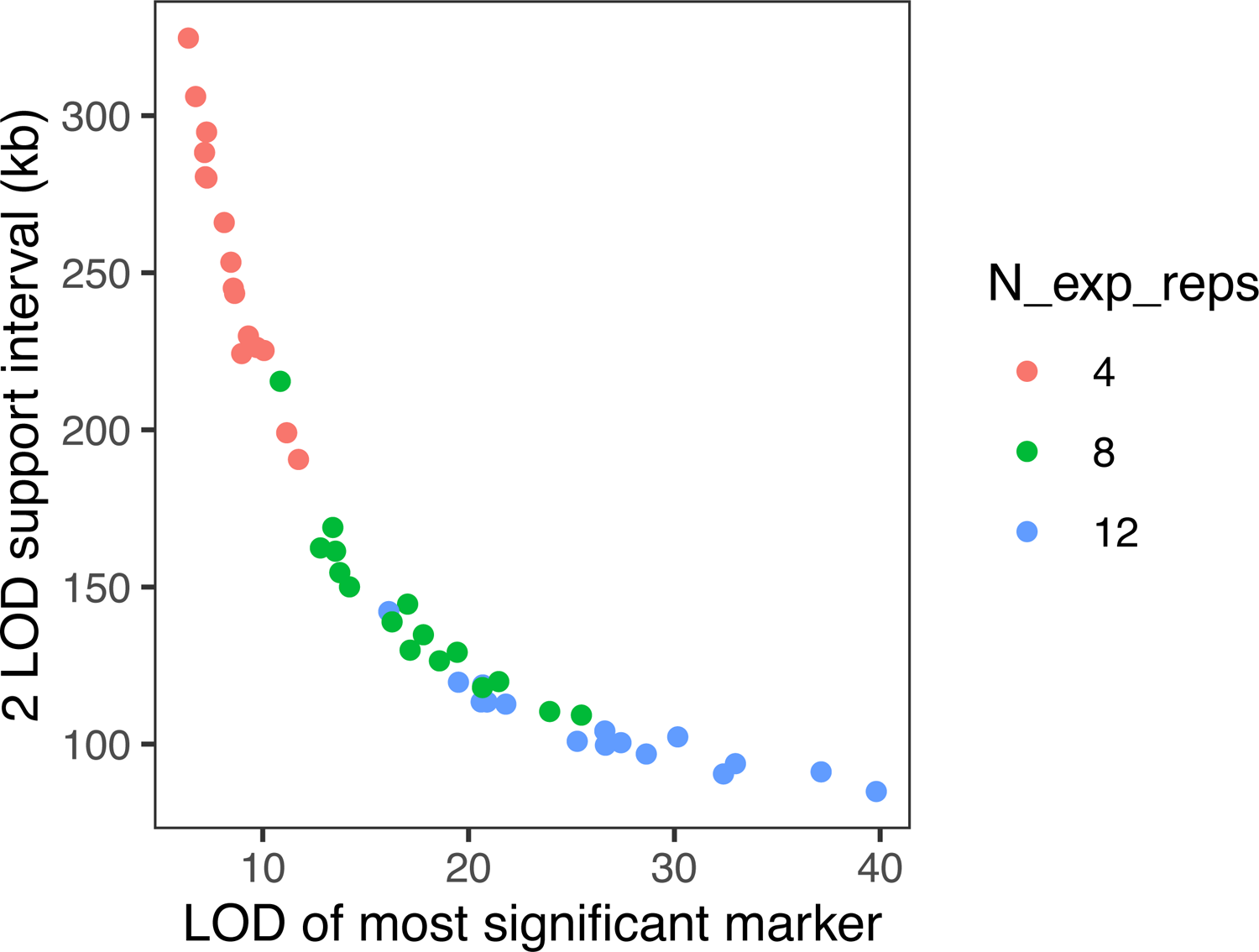
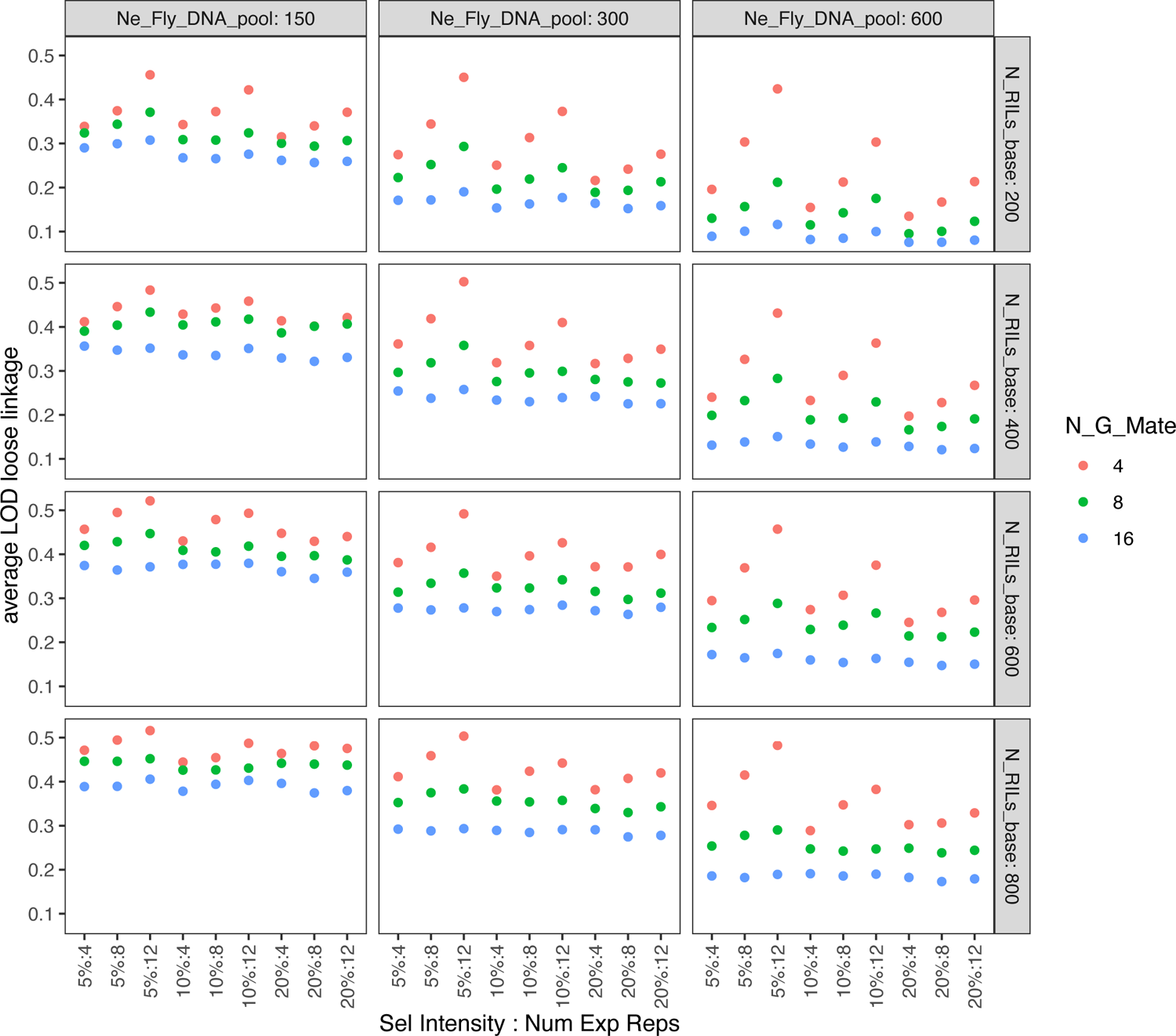
(A) QTL localization for an X-QTL experiment with a single replicate. All parameters identical to Figure 3, except we only present a single realization of the X-QTL experiment. This figure highlights the strong correlation in test statistics at adjacent markers, indicating that we are employing a sufficient marker density - 200 markers along a single chromosome arm (3L) - in our QTL localization simulations. **(B) Average 2-LOD support interval as a function of LOD score at the most significant marker.** Experiment simulates four generations of random mating following base population establishment for different numbers of experimental replicates. Different points are different combinations of design parameters (*N_DNA_* = 300, 600; *N_RIL_* = 200, 400, 600, 800; *i* = 5%, 10%). **(C) Average LOD score at markers loosely linked to causative region.** Data suggests some inflation of LOD scores as a function of the number of generations of random mating following base population establishment, *N_DNA_*, *N_RIL_*, *i*, and the number of experimental replicates.

**Figure S3:**
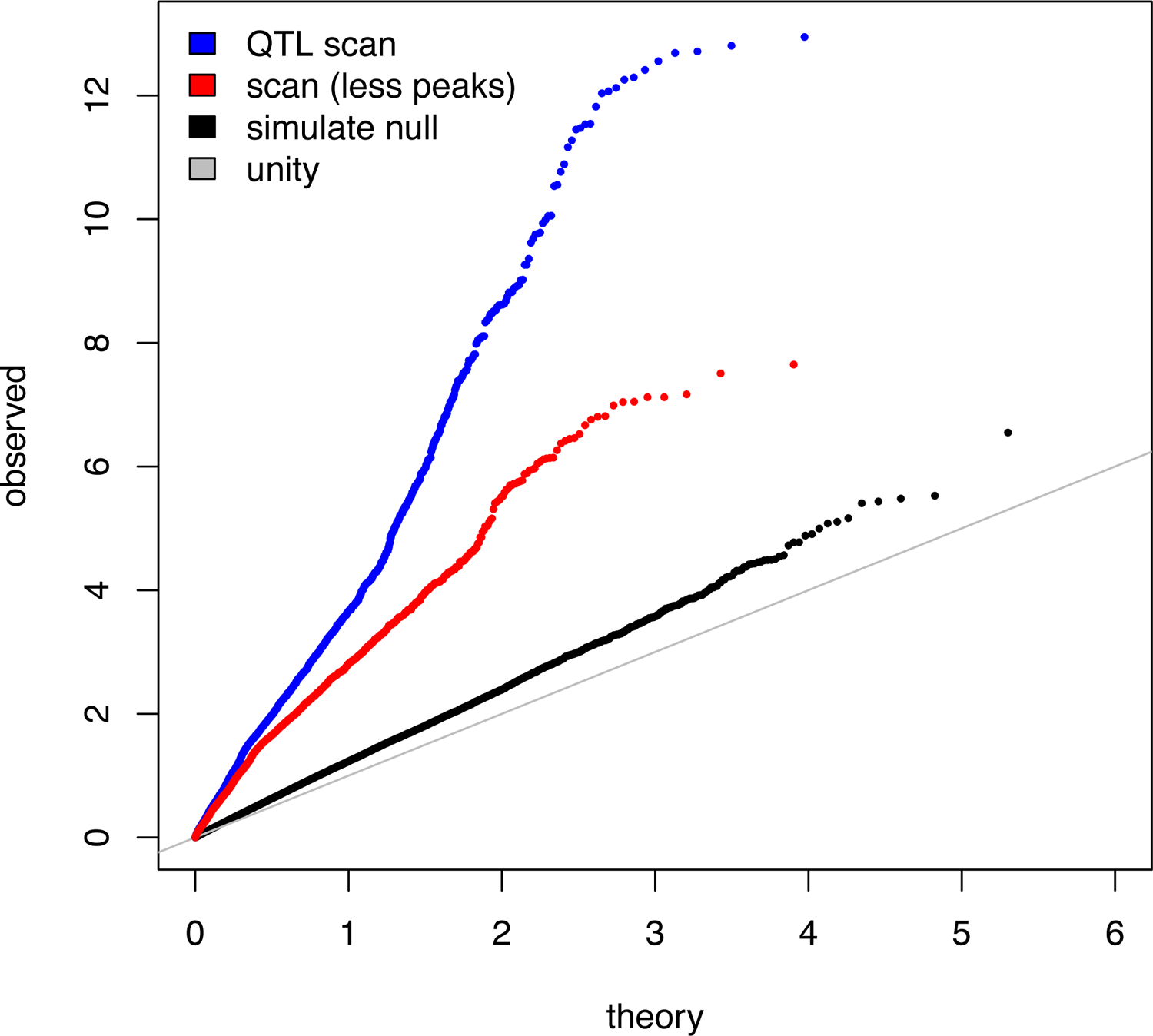
QQ-plots: Theoretical −log_10_(*p*) quantiles on the x-axis, and simulated quantiles (black) or observed quantiles (red and blue) from the caffeine resistance mapping experiment on the y-axis. The grey line is unity, and would be expected if observed quantiles matched theory under a model with no QTL. The black points are observed quantiles under a simulated experiment comparing two control draws of the same size from the base population of size *N_DNA_* = 150 from a 4-fold replicated simulated experiment. There is a slight inflation of the test statistic due to sampling variation. The blue points are observed quantiles under the empirical full coverage X-QTL scan for caffeine resistance loci, and red points are observed quantiles from the same scan with 2-Mb intervals centered on the seven detected peaks (Table 1) removed.

**Figure S4:**
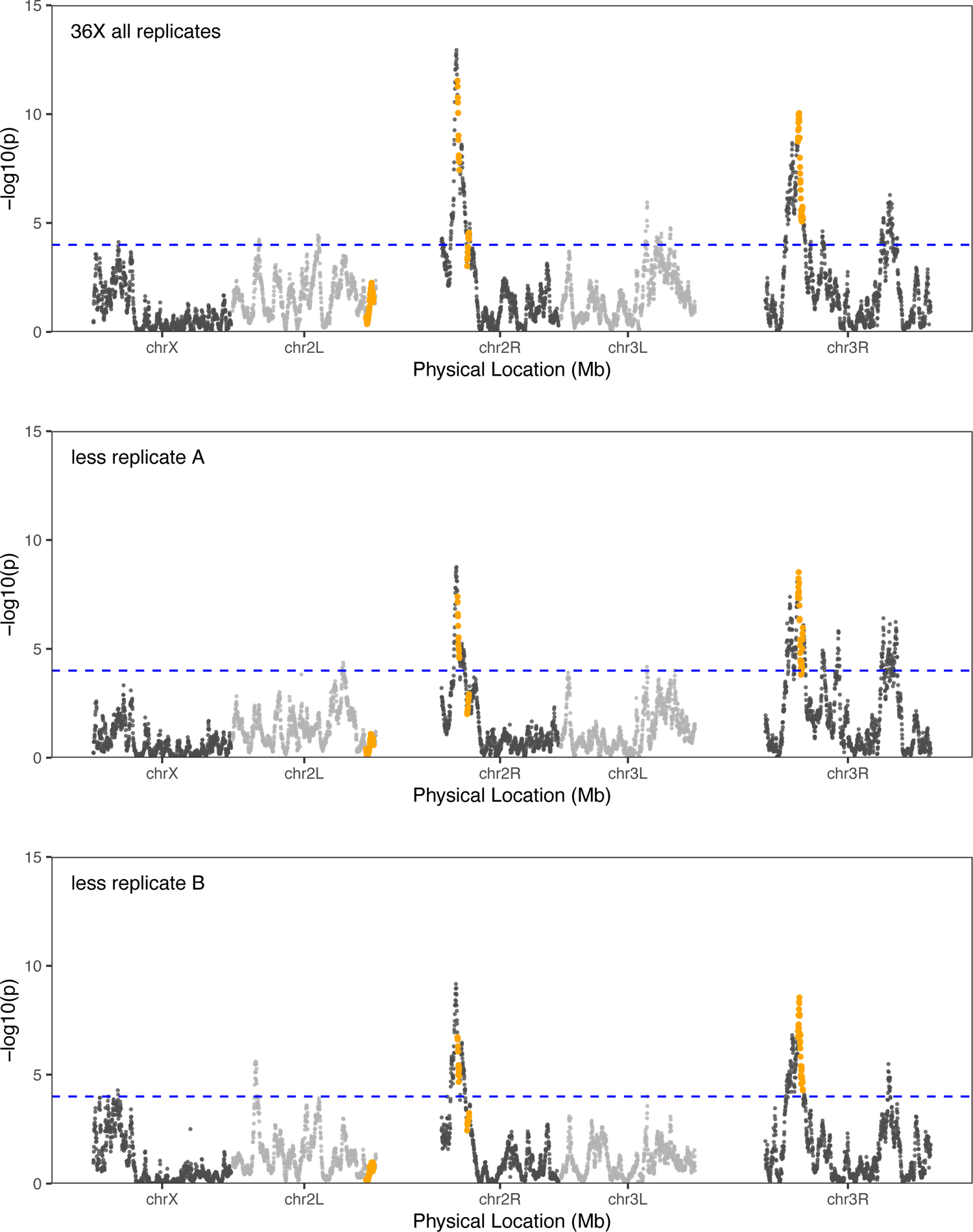
**A QTL scan at ∼36X coverage with some replicates dropped**. The upper panel is for the entire dataset (all 4 replicates), while the two lower panels drop either replicate “A” (where a fraction of the animals were dispensed into phenotyping assay tubes automatically) or replicate “B” (where, similarly to replicates “C” and “D”, all animals were dispensed via manual aspiration). The similarity of the LOD profiles in the lower two panels implies that the method of dispensing does not markedly contribute to the outcome.

**Figure S5:**
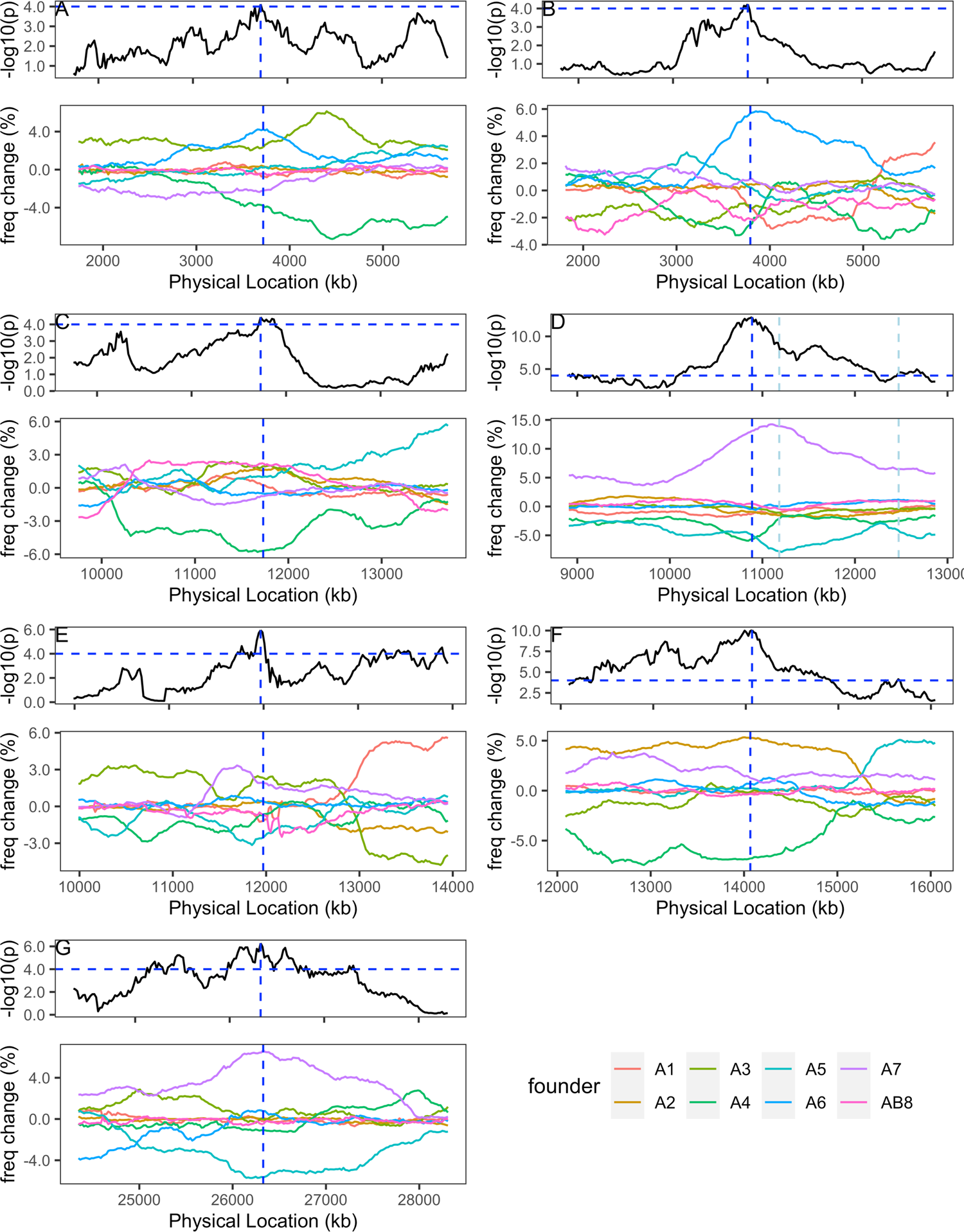
Founder haplotype frequencies at caffeine X-QTL. This figure is identical to Figure 5 except the LOD scores and frequencies are calculated from the lower coverage, ∼35X, sequencing data.

**Figure S6:**
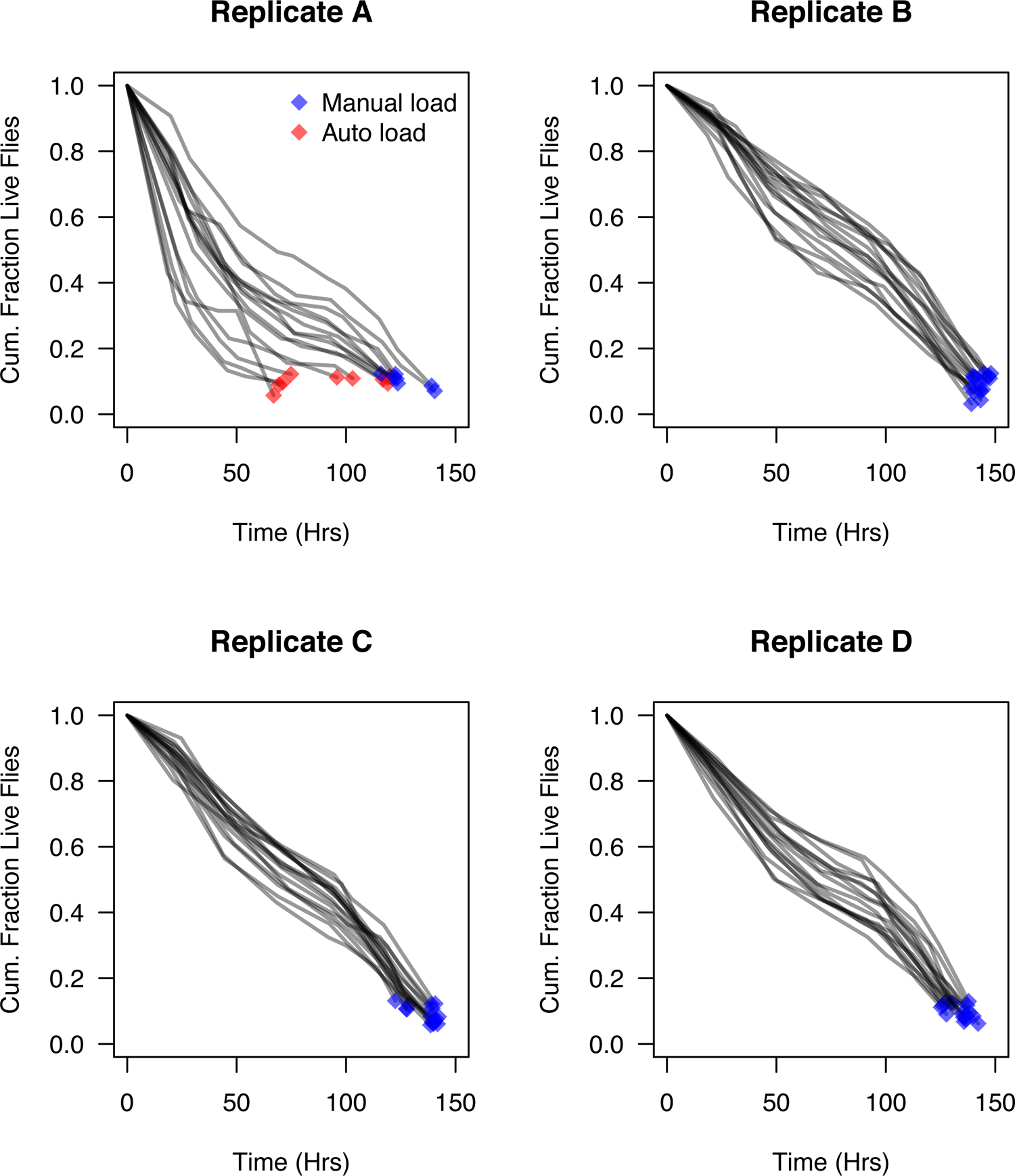
Mortality trajectory of flies from each replicate. Each curve in each replicate-specific plot represents the fraction of flies alive per tray at a series of timepoints throughout the experiment. Trays typically contained 160 flies at the start of each replicate experiment, each fly held singly in an activity monitor tube. Dead flies were typically counted twice per day, and the mortality trajectory is roughly consistent across trays. For replicate A, a fraction of the flies were automatically loaded into monitor tubes (red) and a fraction of the flies were manually loaded (blue), as they were for replicates B-D. Automatic loading led to a reduced lifespan on average, although this variation in experimental method appears not to have had a dramatic effect on the mapping results (see Figure S4).

**Table S1.** Details of each replicate caffeine resistance experiment using the mixed population of DSPR pA RILs.

| Activity | Replicate A | Replicate B | Replicate C | Replicate D |
| --- | --- | --- | --- | --- |
| Generations since population founding | 1 | 3 | 4 | 5 |
| Egg collection date <sup>a</sup> | 25-26/7/19 | 8-9/8/19 | 30-31/8/19 | 5-6/9/19 |
| Number of vials of eggs collected | 105 | 73 | 84 | 54 |
| First flies evident in rearing vials | 3/8/19 | 17-18/8/19 | 7-8/9/19 | 14-15/9/19 |
| Transfer adults from each rearing vial to a mixed-sex housing vial | 5/8/19 | 19/8/19 | 9/9/19 | 16/9/19 |
| Experimental female collection date | 6/8/19 | 20/8/19 | 10/9/19 | 17/9/19 |
| Housing of experimental females | 1 vial of 30 per rearing vial | 2 vials of 22 per rearing vial | 2 vials of 19 per rearing vial | 2 vials of 28 per rearing vial |
| Date flies loaded into activity monitor tubes | 8/8/19 | 22/8/19 | 12/9/2019 | 19/9/19 |
| Estimated age of experimental females on experiment start | 3-5 days old | 3-5 days old | 3-5 days old | 3-5 days old |
| Total number of control flies collected | 250 | 250 | 250 | 250 |
| Total number of flies assayed | 2,337 | 2,572 | 2,535 | 2,563 |
| Total number of selected flies collected | 241<br>(10.3 %) | 235<br>(9.1 %) | 228<br>(9.0 %) | 254<br>(9.9 %) |

<sup>a</sup> day\_start-day\_end/month/year

**Table S2.**
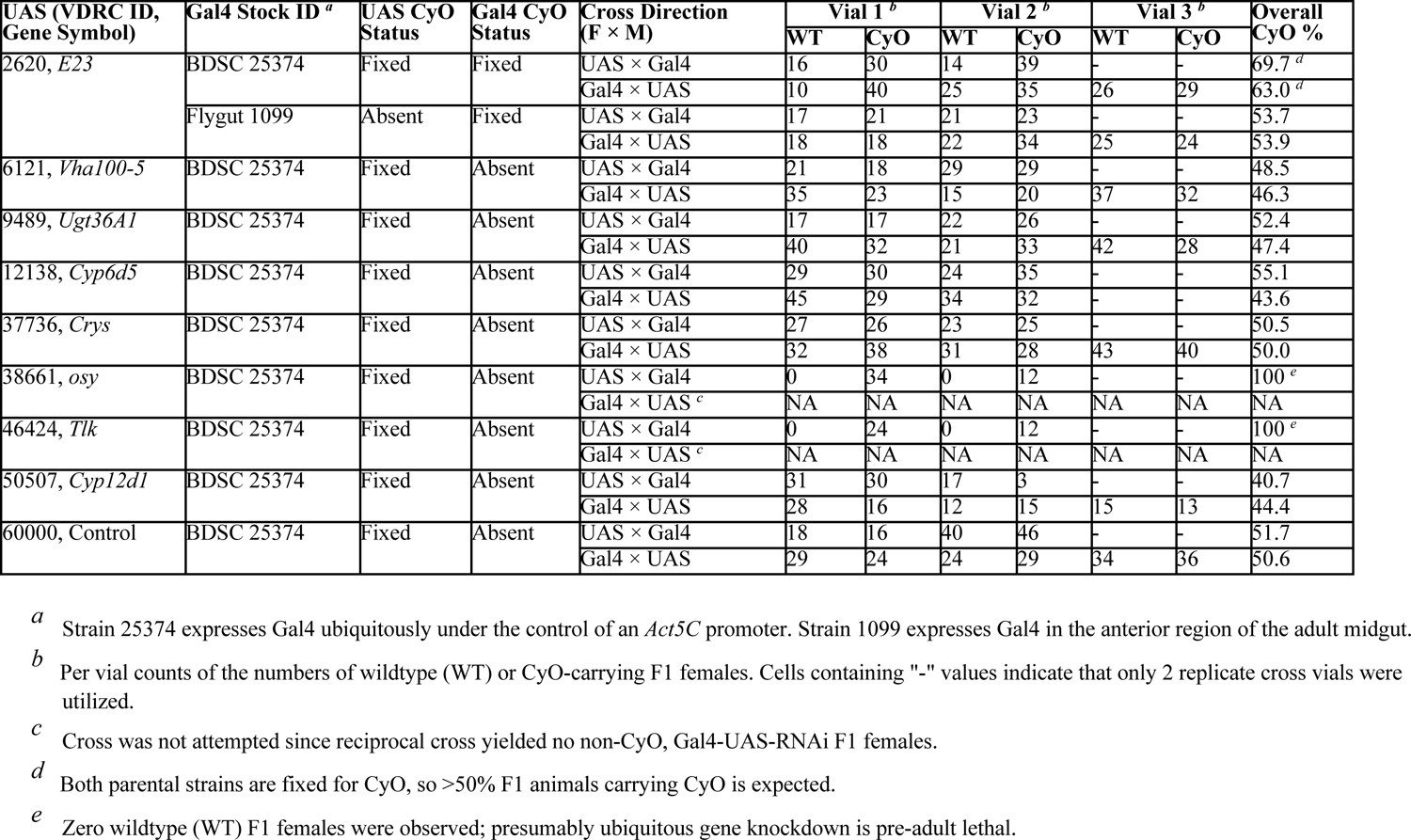
Counts of CyO balancer-containing animals per Gal4 × UAS cross.

**Table S3.** D. melanogaster release 6 euchromatic boundaries

| Arm | Left boundary | Right boundary |
| --- | --- | --- |
| X | 277,911 | 18,930,000 |
| 2L | 82,455 | 19,570,000 |
| 2R | 8,860,000 | 24,684,540 |
| 3L | 158,639 | 18,438,500 |
| 3R | 9,497,000 | 31,845,060 |

## SUPPLEMENTARY TEXT

**Text S1.** Media recipes employed in the study.

### APPLE JUICE AGAR

For 1-liter of apple juice agar:

750-ml water

20-g agar (Genesee Scientific; 66-111)

Mix using stirring hotplate until mix boils

250-ml apple juice (store bought) 25-g sugar (store bought)

Mix in a beaker

Add to boiling water/agar mix

Lower temperature and continue to heat/stir for ∼15-min

Remove from stirring hotplate to orbital shaker to cool

5-ml 95% ethanol

1.5-g tegosept (Genesee Scientific; 20-258)

Dissolve tegosept in ethanol in 50-ml centrifuge tube

Add to water/agar/juice/sugar mix when it has reached ∼60°C

Pour into petri dishes

Avoid generating bubbles, but if some form, use bunsen burner flame to remove them

### LIVE YEAST PASTE

Mix approximately equal volumes of water and active dry yeast (Genesee Scientific; 62-103) until it is smooth, and achieves the consistency of toothpaste.

### CORNMEAL-YEAST-MOLASSES REARING/HOUSING MEDIA

28.5-liters water

280-g agar (Genesee Scientific; 66-111. This is based on a gel strength of 1,090 g/cm^2^, and will change depending the batch)

Add water to steam kettle, turn on electric mixer, and slowly add agar

Bring mix to a boil

3,200-ml molasses (Genesee Scientific; 62-117)

Reduce the kettle pressure to reduce the heat slightly

Add molasses, and bring mix back to a boil

4-liters water

1,460-g inactive dry yeast (Genesee Scientific; 62-107)

Mix in bucket using paint-stirring drill attachment

4-liters water

2,600-g yellow cornmeal (Genesee Scientific; 62-101)

Mix in bucket using paint-stirring drill attachment

Add both the water/yeast and water/cornmeal mixes to the steam kettle

Bring mix back to boil, and simmer for ∼15-min

Release pressure from steam kettle, but continue to stir with electric mixer

330-ml water

259-ml propionic acid (ThermoFisher; A258-500)

31-ml phosphoric acid (85%; ThermoFisher; A242-500) Pour mix into kettle

400-ml 95% ethanol

1.5-g tegosept (Genesee Scientific; 20-258) Dissolve tegosept in ethanol

Pour mix into kettle

Fill vials/bottles

### CORNMEAL-YEAST-DEXTROSE ASSAY MEDIA

For ∼1.5-liters of media:

1,028-ml water

7.5-g agar (Genesee Scientific; 66-111)

Mix using stirring hotplate until mix boils

Reduce heat and continue boiling for ∼15-min until mix is clear

180-ml water

45-g inactive dry yeast (Genesee Scientific; 62-107)

81-g yellow cornmeal (Genesee Scientific; 62-101)

96-g dextrose (Fisher Scientific; D16-1)

Mix in a beaker

Stir into agar/water mix

Boil for ∼10-min (manually stir frequently to avoid burning)

Remove from stirring hotplate to orbital shaker to cool

18-ml acid mix (see below*)

30-ml tegosept/ethanol mix (see below**)

Add to water/agar/yeast/cornmeal/dextrose mix when it is ∼65°C

Move 1-liter of media to fresh beaker

10-g caffeine (SigmaAldrich, C0750)

Add to mix when it is ∼55°C

* Acid mix:

418-ml propionic acid (ThermoFisher; A258-500), 50-ml phosphoric acid (85%; ThermoFisher; A242-500), 532-ml water

** Tegosept / ethanol mix:

3-g tegosept (Genesee Scientific; 20-258) dissolved in 30-ml 95% ethanol

**Text S2.** Pooled fly DNA isolation protocol.

1. Homogenize pool of ∼250 flies in 2-ml of 1X PBS using glass beads with a Mini-BeadBeater-96 (Biospec).
2. Add 4-ml of cold cell lysis buffer* to sample, and subject to 3-4 strokes of both the “loose” and “tight” pestles of a glass dounce tissue grinder (Wheaton, 7-ml).
3. Using a wide-bore pipet tip, move 600-ul of the resulting slurry to a 1.7-ml microcentrifuge tube, incubate at 65°C for 25-min, and cool to room temperature.
4. Add 3-μl of RNase A solution* to lysate, mix the tube by inverting 25 times, incubate at 37°C for 40-min, and rapidly cool to room temperature by placing sample on ice.
5. Add 200-μl of protein precipitation solution* to the lysate, vortex on high speed for 20 seconds, place sample on ice for 5-min, and centrifuge at 14,000-rpm** for 3-min.
6. Move supernatant to new 1.7-ml microcentrifuge tube, and centrifuge at 14,000-rpm** for 1-min.
7. Move supernatant to a 1.7-ml microcentrifuge tube containing 600-μl of isopropanol (avoiding any remaining detritus), mix tube by inverting 50 times, centrifuge at 14,000-rpm** for 1-min, and gently pour off supernatant.
8. Add 600-μl of 70% ethanol, invert tube 2-3 times to wash pellet, centrifuge at 14,000-rpm** for 1-min, pipette off supernatant, leave tube inverted to air dry for 15-min, and resuspend with 50-μl of Qiagen EB buffer***.

* Cell lysis buffer, RNase A solution, and protein precipitation solution are part of the Gentra Puregene Cell Kit (Qiagen, 158767).

** This is >20,000-g (rcf) in our Eppendorf instrument.

*** Qiagen, 19086.

